# ILP4 and InR regulate Paclitaxel-induced hypersensitivity differently in *Drosophila* larvae

**DOI:** 10.1101/2025.09.12.675934

**Authors:** Sreepradha Sridharan, Yogesh Srivastava, Ashleigh Ogg, Yan Wang, Michael J. Galko

## Abstract

Paclitaxel (PTX), a chemotherapeutic that stabilizes microtubules, induces nociceptive hypersensitivity and sensory neuron damage in humans, mice, and flies. To enhance our basic understanding of PTX-induced effects we undertook a molecular/genetic dissection of PTX-induced nociceptive hypersensitivity. Larvae fed viable doses of PTX exhibited dose-dependent hypersensitivity to sub-noxious thermal stimuli. Hypersensitivity developed rapidly and did not completely resolve at the larval stage. Live imaging of peripheral thermal nociceptors showed that lower doses of PTX (< 10 µM) caused hyper-sprouting of tertiary dendritic branches. At 10 µM and above, dendritic beading was observed. PTX-induced hypersensitivity does not depend on signaling pathways previously implicated in acute injury-induced nociceptive sensitization. However, the insulin-like peptide 4 (ILP4), was required for PTX-induced thermal hypersensitivity at 10 µM PTX. Surprisingly, RNAi targeting the insulin receptor (InR) in nociceptors increased PTX-induced hypersensitivity, suggesting that ILP4 does not activate InR in this context. The salivary gland is likely the primary tissue source of functional ILP4. ILP4 mutant larvae did not exhibit PTX-induced beading (10 µM) but did exhibit hypersprouting at lower PTX concentrations. In summary, our model of PTX-induced hypersensitivity reveals a disconnect between hypersensitivity and neuronal morphology and a genetic separation of ILP4 and InR in PTX-induced hypersensitivity.

## Introduction

Paclitaxel (PTX) is a common chemotherapeutic agent used for treating ovarian, breast, and non-small cell lung cancer. The taxane PTX binds tubulin (Gallego-Jara *et al*., 2020) and effectively arrests mitosis by preventing microtubule depolymerization during prometaphase (Weaver, 2014). Nearly 97% of patients who receive PTX experience pain hypersensitivity or other symptoms of chemotherapy-induced peripheral neuropathy (Klein and Lehmann, 2021). Clinical survey of patient symptoms showed that 88% of patients felt acute pain symptoms within four days of starting treatment (Reeves *et al*., 2012). *In vitro* studies of PTX suggest that pain hypersensitivity may be caused by several mechanisms including impaired axonal transport (Theiss and Meller, 2000) related to alterations in microtubule polarity (Shemesh and Spira, 2010). Cultured rat sensory neurons exhibit increased spontaneous activity on PTX treatment (Zhang and Dougherty, 2014) a response that is consistent with in vivo hypersensitivity. Skin biopsies from patients receiving docetaxel suggest that degradation occurs in small nerve fibers and correlates with pain hypersensitivity and alterations to nerve conduction (Kroigard *et al*., 2014).

Studies in mice have shown that PTX causes thermal hyperalgesia (increased pain to noxious stimuli), mechanical and chemical allodynia (increased pain behavior in response to previously non-noxious stimuli), and loss of sensitivity to chemical stimuli (Masocha, 2014; Toma *et al*., 2017). Morphologically, PTX administration in mice causes retraction and loss of intra-epidermal nerve fibers (Krukowski *et al*., 2015; Toma *et al*., 2017) similar to the dying-back of axons seen in patients. Similar morphological and behavior effects have been observed in non-mammalian vertebrates like Zebrafish (Lisse *et al*., 2016). In vertebrate models of PTX-induced hypersensitivity the relationship between hypersensitivity and morphological changes in sensory neurons remains unclear.

The combination of genetic tools and established nociception assays make *Drosophila* larvae a powerful system for advancing our understanding of nociception, nociceptive sensitization, and their underlying mechanisms (Im and Galko, 2012). *Drosophila* have dedicated multidendridic nociceptors that can be subdivided into four classes (Grueber *et al*., 2002). The class IV sensory neurons are responsible for detection of noxious thermal and mechanical stimuli (Tracey *et al*., 2003; Hwang *et al*., 2007) and optogenetic activation of these neurons causes aversive rolling. (Hwang *et al*., 2007; Turner *et al*., 2016). As in vertebrates, conserved transient receptor potential ion channels like painless (Tracey *et al*., 2003) and TRPA1 (Neely *et al*., 2011; Zhong *et al*., 2012) mediate baseline nociceptive responses. There is a dedicated neuronal circuitry in fly larvae that includes second-order and higher-order central neurons that process nociceptive information (Boivin *et al*., 2023) and the baseline nociceptive response can be sensitized by injury (Babcock *et al*., 2009; Khuong *et al*., 2019) and even disease states like diabetes (Im *et al*., 2018).

Exposure of *Drosophila* larvae to vinca alkaloid chemotherapeutics causes both mechanical and thermal hypersensitivity (Boiko *et al*., 2017). Studies of PTX-induced neuropathy using *Drosophila* have mainly focused on neuron degeneration (Bhattacharya *et al*., 2012; Brazill *et al*., 2018; Kim *et al*., 2020; Shin *et al*., 2021). Larvae fed PTX have increased dendritic branching density (Brazill *et al*., 2018; Shin *et al*., 2021) as well as axonal degradation (Bhattacharya *et al*., 2012). Behaviorally, larvae also show increased hypersensitivity to thermal stimuli (Brazill *et al*., 2018). Whether non-toxic (no neuronal degradation) concentrations of PTX cause hypersensitivity and whether this hypersensitivity is similar genetically to injury- or disease-induced nociceptive hypersensitivity remain open questions in the field.

In *Drosophila*, the insulin receptor (InR) mediates the persistence of nociceptive hypersensitivity after UV injury and in diabetic-like states. The latter are of particular interest to PTX-induced hypersensitivity because diabetic patients (Lauria *et al*., 2003), mice (Toma *et al*., 2017), and flies (Im *et al*., 2018) all experience nociceptive hypersensitivity that can be accompanied by damage to peripheral neurons (Feldman *et al*., 2019). *Drosophila* have eight insulin-like peptides (ILPs) and one insulin receptor, InR (Brogiolo *et al*., 2001; Semaniuk *et al*., 2021). Whether ILPs and InR regulate PTX-induced hypersensitivity has not previously been examined.

Here, using a refined model of PTX-induced hypersensitivity we characterize the effects of low concentrations of PTX, the onset and resolution of PTX-induced hypersensitivity, and the relation of hypersensitivity to neuronal morphology. We also identify new molecular regulators of PTX-induced hypersensitivity‒ an ILP (ILP4) and InR‒, as well as identify the functional source of ILP4 peptide for regulating PTX-induced hypersensitivity.

## Materials and Methods

### Fly genetics

Stocks were obtained from the Bloomington *Drosophila* Stock Center, the Vienna *Drosophila* Resource Center, and the Korea *Drosophila* Resource Center. *Drosophila* were reared on standard cornmeal medium under a 12 h light-dark cycle. A table (Table S2) of all genotypes used in the figures in the paper is included in the supplemental material. All crosses were cultured at 25 °C. *w^1118^* was used as a control strain. The *ILP* mutants (ILP2, ILP3, ILP4, ILP5, ILP6, ILP7, and ILP8) used (Gronke *et al*., 2010; Colombani *et al*., 2012; Garelli *et al*., 2012) were tested as homozygotes.

The Gal4/UAS system (Brand and Perrimon, 1993) was used to direct tissue-specific expression of various transgenes. UAS-lines used: *UAS-wengen^IR^*(here named *UAS-TNFR^RNAi^*) was used to reduce TNF signaling (Kanda *et al*., 2002; Babcock *et al*., 2009). *UAS-Tachykinin^RNAi^*(Dietzl *et al*., 2007; Im *et al*., 2015) and *UAS-Smo^RNAi^* (Dietzl *et al*., 2007; Babcock *et al*., 2011) were used to reduce tachykinin signaling, and hedgehog signaling, respectively. As previously, *UAS-InR^RNAi^* was used to reduce insulin signaling (Perkins *et al*., 2015; Im *et al*., 2018). *UAS-mCD8-GFP* (Lee and Luo, 1999) and *UAS-DsRed2-Nuc* (Lesch *et al*., 2010) were used in combination with various Gal4 drivers (see below) to visualize class IV nociceptive sensory neurons and ILP4 expressing tissues. *UAS-ILP4* (Ikeya *et al*., 2002) was used for tissue-specific rescue experiments. Four different *UAS-ILP4^RNAi^* lines (BL43288, BL31377, VDRC 105566, VDRC 12890) (Dietzl *et al*., 2007; Ni *et al*., 2011) were tested for efficacy with *Da-gal4*, (Wodarz *et al*., 1995) which expresses ubiquitously. *UAS-ILP4 RNAi* line #12890 was then used to knock down ILP4 in each tissue that expresses it.

Tissue-specific Gal4 drivers used: To drive expression in nociceptive class IV sensory neurons we used *ppk1.9-gal4* (Ainsley *et al*., 2003). We used *btl-gal4* (Shiga *et al*., 1996) to drive expression in trachea*, gr66a-Gal4* (Dunipace *et al*., 2001) to drive expression in the dorsal organ/terminal organ (DO/TO). *Sage-gal4* (Fox *et al*., 2013) and *Fkh-Gal4* (Henderson and Andrew, 2000) were used to drive expression in salivary glands. *ILP4-gal4* (Min *et al*., 2016) was used to drive expression in all ILP4-expressing tissues. *Ppk1.9-Gal4 > UAS-mCD8-GFP* was used to assess neuronal morphology in otherwise wildtype larva as in (Lopez-Bellido *et al*., 2019) while *Ppk1.9-Gal4 > UAS-mCD8-GFP; ILP4-mut* was used to assess neuronal morphology of larva lacking ILP4.

### PTX feeding and survival experiments

Solid PTX (ThermoFisher #P3456) was resuspended in DMSO (Fisher Chemical #BP 231-1) to a concentration of 10 mM and this master stock solution was stored at -20 °C until use. The PTX master stock was further diluted in deionized water for use in experiments. These dilutions were based upon the toxicity (survival) and behavioral potency of the particular batch of PTX used. For nearly all experiments, the final standard assay concentration was 10 µM PTX-meaning this was the highest concentration that did not adversely affect survival at the larval stage. However, the PTX batch that was used for the salivary gland rescue experiment (Figure 5L) was more potent and was used at 6 µM, the highest concentration of this batch that did not adversely affect survival and largely matched the behavioral potency of previous batches at 10 µM. Instant fly food containing blue dye (Carolina Biological #173210) was mixed with PTX and deionized water. For survival experiments, 0.25 g of instant fly food was mixed with a total volume of 1 mL deionized water containing PTX or DMSO vehicle (control). This mixture was placed in 1.5 mL clear dram vials (Fisher Brand#03-339-26B) and 20 mid-third instar larvae of both sexes were placed on the surface of the food and incubated at 25 °C. Larvae remained in food until pupariation. Pupae that formed from PTX-fed larvae were counted and compared to their respective DMSO controls using unpaired t-tests.

### Thermal nociception assay and quantitation of behavioral data

1.25 g of instant fly food was mixed with a 5 mL volume of PTX and deionized water and placed in *Drosophila* vials (Genesee Scientific, #32-110). 30-40 larvae were added to the food per vial and incubated at 25 °C. Thermal nociceptive assays were performed at room temperature using a custom-built metal probe (ProDev Engineering) with a thermal control unit as previously reported (Babcock *et al*., 2009). Larvae were assayed at a probe temperature of 38.5°C unless otherwise indicated and the probe contacted the dorsal aspect of larval segment A4-A6. We scored the noxious withdrawal response of aversive rolling for up to 20 s of probe contact. Only larvae that exhibited a full 360° body roll were scored as responders and the latency from initial probe contact rounded to the nearest second, was recorded. Data was plotted as cumulative percent response versus latency and statistical significance was determined using the Log-rank Mantel-Cox test, comparing two groups (control versus experimental) at a time. Because this behavioral assay varies slightly with the particular probe and experimenter, the dose-response curve of rolling behavior versus set temperature for this study is reported in Figure S1.

The behavioral response differences between larvae of the same genotype +/-PTX (as in Figure S5) were assessed by comparing the average latency gap between the two datasets (-PTX versus + PTX). As above, a latency of 20 s was defined as a non-responder (NR) so the maximum response latency = 19 s and the maximum latency gap = 18 s (a responder of 19s versus a responder of 1s). All pairwise combinations between untreated (A) and treated (B) groups were computed in either direction as follows: AB (A − B) and BA (B − A). So, with datasets of 90 larvae, there are 90 x 90 = 8100 latency gap comparisons. Also note that with this analysis it is possible to have negative values. If either A or B was a non-responding larva, the calculation was labeled as “noGap” as follows:

If A = 20 or B = 20 → then value = 0 = “noGap”

And: If A < 20 and B < 20 → then value = 1 = “a latency gap is present”

Comparisons with value = 0 (‘no gap’) were retained in the dataset and contributed to the total number of pairwise comparisons. These were included in the denominator when calculating cumulative response percentages as follows: Cumulative % response = (number of comparisons of value = 1/total number of comparisons) X 100. What is achieved by this analysis is a computation of the “average latency gap” observed for larvae of the same genotype +/-PTX. Basically, two lifespan curves (genotype X + PTX and genotype X – PTX) are converted into a single curve (with possible positive and negative values) that reflects the average effect of the drug. The resulting comparisons between two different genotypes were statistically compared using a pairwise log-rank test. For each genotype and orientation (AB or BA), survival-type curves of cumulative response percentage as a function of latency gap magnitude using basic R’s lm() function were compared. All analyses and visualizations were conducted in R (version ≥ 4.0.0) using the tidyverse, ggplot2, readxl, and broom packages.

### Microscopy to assess nociceptive sensory neuron morphology

Third instar larvae of the genotype *ppk1.9-gal4>UAS-mCD8-GFP/CyO* or *Ppk1.9-Gal4 > UAS-mCD8-GFP/CyO; ILP4-mut* were anesthetized using anhydrous ethyl ether (Fisher Chemical, #E138-500) and mounted live using halocarbon oil (Millipore Sigma, #H8898) and ether according to previously published protocol (Das *et al*., 2017; Lopez-Bellido and Galko, 2020). Fluorescent class IV peripheral sensory neurons were imaged by confocal fluorescence microscopy using a Nikon Eclipse Ti equipped with a Nikon GaAsP detector. The laser used was 487 nm with emission filter wavelengths 500-550nm. A Nikon Plan Apochromat VC 20x/0.75NA objective was used to capture images. All images were acquired with NIS Elements and reported as maximum intensity projections.

### Microscopy to assess ILP4 expression

Third instar larvae of the genotype *ILP4-Gal4 > UAS-DsRed2Nuc* were anesthetized using anhydrous ethyl ether (Fisher Chemical, #E138-500) and mounted live using halocarbon oil (Millipore Sigma, #H8898) and ether according to previously published protocol (Das *et al*., 2017; Lopez-Bellido and Galko, 2020). For the salivary glands, which is a more internal tissue not easily visualized in the live preparation, a dissected wholemount preparation was used. Briefly, the head of each larva was removed using forceps as shown in (Cai *et al*., 2010). Once removed, larval heads with salivary glands attached were placed into a 1.5mL microcentrifuge tube filled with 1X Phosphate-buffered saline (PBS). Larval samples were fixed with 3.7% formaldehyde on ice for 25 minutes. Samples were washed 4 times with PBST (PBS plus 1 % Tween-20-(Fisher BioReagents, #BP337-500) for 10 minutes each before placing 50 µL Vectashield (Vector Laboratories, #H-1000-10) into the microcentrifuge tube.

Samples were transferred from the microcentrifuge tube to a clean microscope slide. Using two microdissection pins (Fine Science Tools, # 26002-10) affixed to a holder (Fine Science Tools, #26016-12), excess cuticle or fatty tissues (if present) were removed. Samples in ∼9 µL of clean Vectorshield were then transferred to a clean microscope slide. A coverslip (22 mm sq, Corning #2865-22) was then placed over the samples and sealed with clear nail polish.

Mounted tissues were imaged by confocal fluorescence microscopy using a Nikon Eclipse Ti equipped with a Nikon GaAsP detector. The laser used was 561 nm with emission filter wavelengths 571 nm - 617 nm. A Nikon Plan Apochromat VC 20x/0.75NA objective was used to capture images. All images were acquired with NIS Elements and reported as maximum intensity projections.

### Neuronal image analysis

Sholl analysis (Sholl, 1953) of neurons was done in Imaris Microscopy Image Analysis Software (Oxford Instruments, Version 10.2). Parameters were defined using Imaris Filament Tracer. Imaris automatically measured dendrite length, branching density, and performed the Sholl analysis. Sholl circles were at 5 μm intervals radiating from the soma. Dendrites smaller than 1.86 μm in length were excluded as these were more likely to be transient spines and all other threshold settings were kept consistent for each neuron analyzed.

Degradation quantification was performed using ImageJ. For each image, a mask to highlight beads was created. This mask was overlayed on the original image and the number of highlighted points within the set range were manually counted. Major dendrites (primary branches which emerge from the neuronal soma) and branch points (of any branch level-primary, secondary, or tertiary) were excluded from the count as these are bright in both controls and experimental conditions. This focused the analysis on bright spots (beads) that are distant from the soma and located only along the shaft of secondary or tertiary dendritic processes (as opposed to branch points).

### ILP4/ILP5 binding to InR-expressing HEK-293 cells

Human embryonic kidney cells (HEK293; ATCC CRL-1573) were grown in 24-well tissue culture plates using 12 mm glass coverslips that had been pre-treated with 100 µg/ml poly-D-lysine (Sigma-Aldrich; P4832-50ML) in Dulbecco’s Modified Eagle Medium (DMEM; CorningTM DMEM; Fisher Scientific MT10017CV) supplemented with 10% fetal bovine serum (FBS) (Heat-inactivated FBS; Sigma-Aldrich #F4135) and 1x antibiotic-antimycotic solution (Fisher Scientific MT30004CI). An expression plasmid (pcDNA5/FRT-INR) was constructed (outsourced to BioMatik, Inc.) in which the coding region of the InR cDNA was chemically synthesized with a 5’ Not1 restriction site and a 3’ Apa1 restriction site and then cloned into the multiple cloning site of pcDNA5/FRT (ThermoFisher Scientific V601020) using those restriction enzymes. The construct was sequence-verified. Cells at 50–60% confluence were co-transfected for 48 hrs with 1 µg of exogenous DNA (e.g., pcDNA5/FRT/TO-GFP or pcDNA5/FRT-INR) using Lipofectamine 2000 (Life Technologies 11668019) according to the manufacturer’s instructions. At the end of the transfection period, cells in each well treated for 30 min with 10 µM of the cell-permeable dynamin inhibitor Dyngo-4a (Selleck Chemicals; #S7163) in FluorBrite^TM^ DMEM (Life Technologies A1896701) to inhibit dynamin-mediated endocytosis and thus maximize receptor presence on the plasma membrane. The following fluorescently labeled peptides were used to assess binding: ILP4 (Chain A: IAHECCKEGCTYDDILDYCA; Chain B: RRKMCGEALIQALDVICVNGFT) (Synthesized by Phoenix Pharmaceuticals Inc. lot #436272) and ILP5 (Chain A: GVVDSCCRKSCSFSTLRAYCDS; Chain B: NSLRACGPALMDMLRVACPNGFNSMFAK) (Phoenix Pharmaceuticals Inc. lot #436210). The peptides were dissolved in 10% DMSO to create 100 µM stock solutions and transfected cells were treated with 1 µM of either peptide for 30 min at 37°C in a CO₂ incubator. Following binding, the cells were fixed in 4% paraformaldehyde and then washed four times for 5 minutes each with PBS. A Nikon AXR Confocal Microscope equipped with an Apo-Plan 60×/1.4 NA oil immersion objective was used for fluorescence imaging. NIS-Elements software and the Nikon AXR GaAsP four-channel detector system were used to illuminate the samples and take digital pictures. Z-stacks were obtained with a pixel resolution of 100 nm and a step size of 0.2 µm.

### qRT-PCR to measure ILP4 expression in fly larvae

The Direct-zol™ RNA MiniPrep Plus kit (Zymo Research, Cat. #R2072) was used to extract total RNA from 15 larvae of the desired genotypes. The concentration and purity of RNA were assessed with a NanoDrop ND-1000 spectrophotometer. First-strand cDNA synthesis was performed using the SuperScript™ IV Reverse Transcriptase kit (Invitrogen, Cat. #18091050) and 1 µg of total RNA template from each sample. To ensure uniformity all samples were diluted and standardized to the same amount (500 ng) before being used in qRT-PCR assays.

PowerUp™ SYBR™ Green Master Mix Kit (Applied Biosystems, Cat. #A25741) was used for the qRT-PCR. Each 20 µL reaction mixture consisted of 10 µL of SYBR master mix, 1 µL of forward primer (ILP4F: 5’-GAGCCTGATTAGACTGGGACTGG-3’ or SRS reference gene: sequence here, 1 µL of reverse primer (ILP4R: 5’-CCCAAACTTACCACTGCTCC-3’ or SRS SRSF:5’-CTGAGAAACGGCTACCACATC-3’ ; SRSR: 5’-ACCAGACTTGCCCTCCAAT-3’ as in (Ponton *et al*., 2011), 1 µL of normalized cDNA (500 ng), and 7 µL of nuclease-free water. For each experimental sample, two technical and three biological replicates were performed. For each primer pair, no-template controls (NTCs) were included. The thermal cycling conditions used on a QuantStudioTM 5 Real-Time PCR System (Applied Biosystems #A28133) included Uracil DNA-glycosylase activation at 50 °C for one min., Dual-Lock Taq polymerase activation at 95 °C for three min, and 40 amplification cycles consisting of denaturation at 95 °C for 30 s, annealing at 60 °C for 30 s, and extension at 72°C for 30 s. The raw Ct results were analyzed in R.

Normalization was performed using the reference gene (SRS; CG8900) signal and ΔCt values were computed by subtracting the average Ct of SRS from the Ct of ILP4 for each sample. The ΔΔCt technique was used to calculate relative expression, using Da-Gal4/+ as the reference genotype. Ct values were normalized to SRS housekeeping gene using the ΔCt or ΔΔCt method using QuantStudio Design & Analysis Software. R (version ≥ 4.0.0) was used for all data processing, statistical analysis, and visualization.

## Results

### Viable doses of PTX cause thermal hypersensitivity

PTX fed to *Drosophila* larvae induces thermal nociceptive hypersensitivity at high concentrations that also adversely affect sensory neuron dendritic morphology (Brazill *et al*., 2018; Shin *et al*., 2021). To test PTX-induced hypersensitivity at lower concentrations that might result in less damage, we first established which doses of PTX were compatible with larval survival to the pupal stage. Second instar larvae were fed PTX ranging from 1 µM to 30 µM, with matched vehicle (DMSO) controls and left to pupariate (Figure 1A and see methods). With most batches of commercially obtained PTX, PTX concentrations above 10 µM significantly decreased larval survival (Figure 1B). For this reason, we only tested larvae at concentrations of 10 µM and below (or the highest dose that did not adversely affect survival-see methods for details) for all further behavioral analysis.

**Figure 1.**
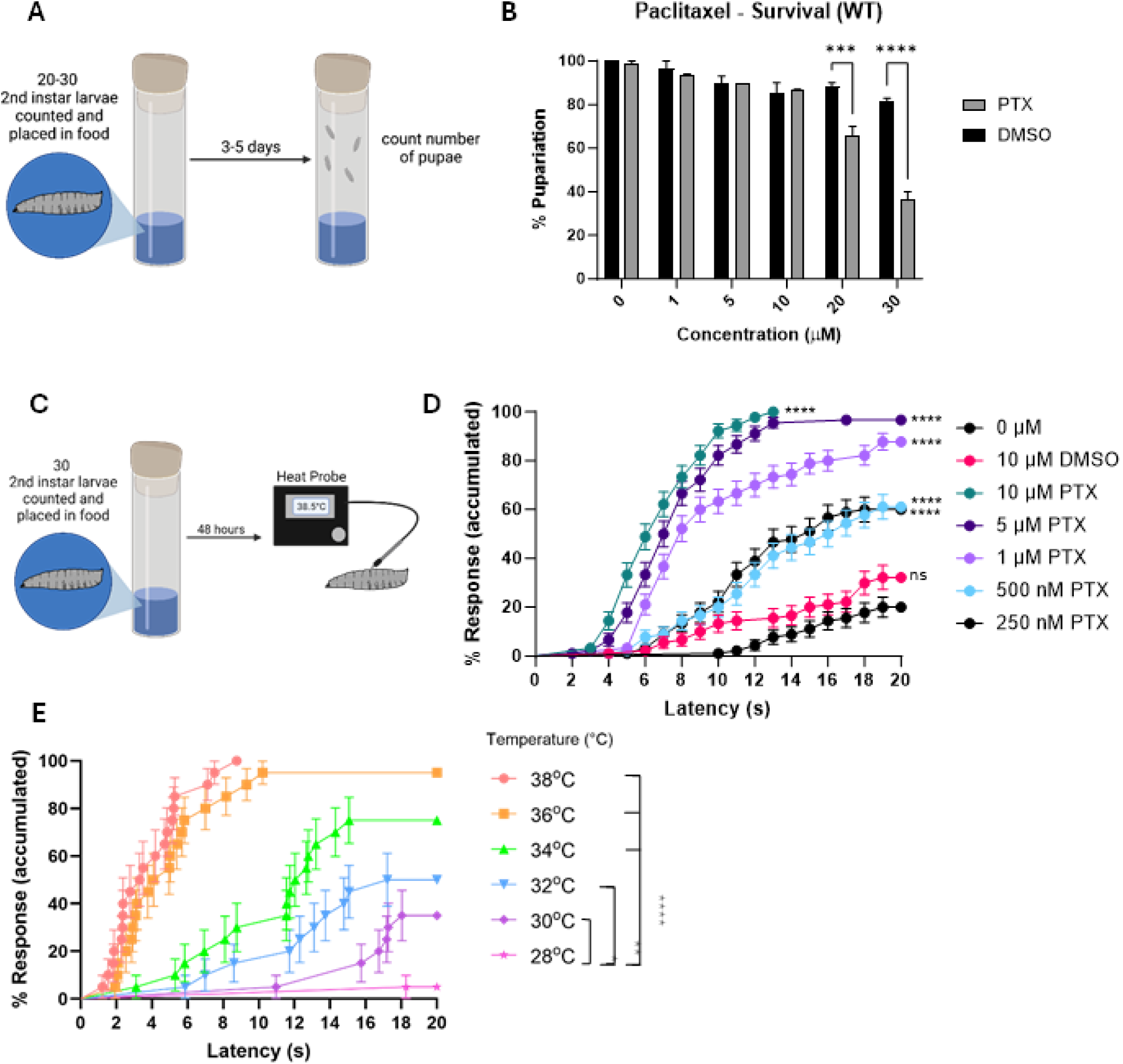
PTX at viable doses causes thermal nociceptive hypersensitivity. A. Cartoon of feeding scheme for assessing survival. B. Quantitation of larval survival to the pupal stage on PTX-containing food versus vehicle only controls. Stats: t-test (n=50, ***p=0.0006, ****p<0.0001(two-way ANOVA)). C. Cartoon of heat probe assay for measuring thermal nociception. D. Dose-response curve of thermal nociceptive hypersensitivity (38.5 °C probe) following 48 hrs feeding on the indicated concentrations of PTX. Data is plotted as cumulative percent of larvae exhibiting aversive rolling behavior versus the latency following probe contact at which rolling was observed (if it was observed within 20 seconds). Controls include larvae fed on normal instant fly food (0 µM) and 10 µM DMSO, the highest concentration of vehicle used. (n = 90 larvae per PTX dose, ****p<0.0001 (Log-Rank Mantel-Cox); ns = not significant) E. Dose response curve of thermal nociceptive hypersensitivity versus probe temperature. Larvae were fed 24 hours on 10 µM PTX and then probed at various temperatures that are normally at or below the border between noxious (∼38 °C) and non-noxious (< 38 °C). Data is plotted as cumulative percent of larvae exhibiting aversive rolling versus the latency following probe contact at which rolling was observed (if under 20 s). Control larvae were fed on normal instant fly food and vehicle (DMSO) without PTX. (n = 30 larvae per setpoint temperature, (Log-Rank Mantel-Cox) comparing each temperature to 28°C as an approximate baseline. (****p<0.0001, **p=0.0011, *p=0.0155).

To assess thermal nociceptive responsiveness, we used a custom-designed heat probe ((Babcock *et al*., 2009) and see methods). Because we were primarily interested in PTX-induced hypersensitivity we assayed larvae at 38.5 °C, a temperature near the non-noxious/noxious threshold (see Figure S1 for dose-response curve of the experimenters and heat probes who performed behavioral analyses in this study). Larvae fed on PTX-containing food for 48 hours were assayed immediately afterward (Figure 1C). Interestingly, larvae exhibited thermal hypersensitivity compared to vehicle controls at PTX concentrations as low as 250 nM (Figure 1D) where over 50 % of larvae respond with aversive rolling. As the concentration of PTX increased the percent of total responders increased, reaching 100 % at 10 µM. Simultaneously, the average response latency decreased with increasing PTX concentration. A second measure of the degree of hypersensitivity caused by PTX feeding was to test larval responsiveness to lower normally non-noxious temperatures. Larvae reared on 10 µM PTX for 24 hours were significantly hypersensitive to the probe all the way down to 32 °C versus vehicle-matched controls (Figure 1E).

### PTX-induced hypersensitivity has a rapid onset and does not resolve at the larval stage

We next sought to better understand how long it takes for PTX-induced hypersensitivity to develop. To study hypersensitivity onset, larvae were fed on PTX-containing food for 2, 4, 8, 16, or 24 hours and compared to larvae fed for 48 hours as above. At 10 µM PTX, larvae fed for only 4 hours had significantly lower hypersensitivity compared to 48 hours (62 % versus 100 %) but were still significantly hypersensitive compared to untreated controls, unlike those fed for only 2 hours (compare 2-hour line in Figure 2A to DMSO control in Figure 1D). Hypersensitivity increased with increasing feeding time until 24 hours, which was not significantly different from a 48-hour feeding. At lower concentrations of PTX (1 and 5 µM), larvae were still hypersensitive within 24 hours however, they had reduced hypersensitivity with 8 hours of feeding (Figure S2). Together, this data shows that onset of PTX-induced thermal hypersensitivity is rapid, even at low concentrations, and peaks by 24 hours of exposure.

**Figure 2.**
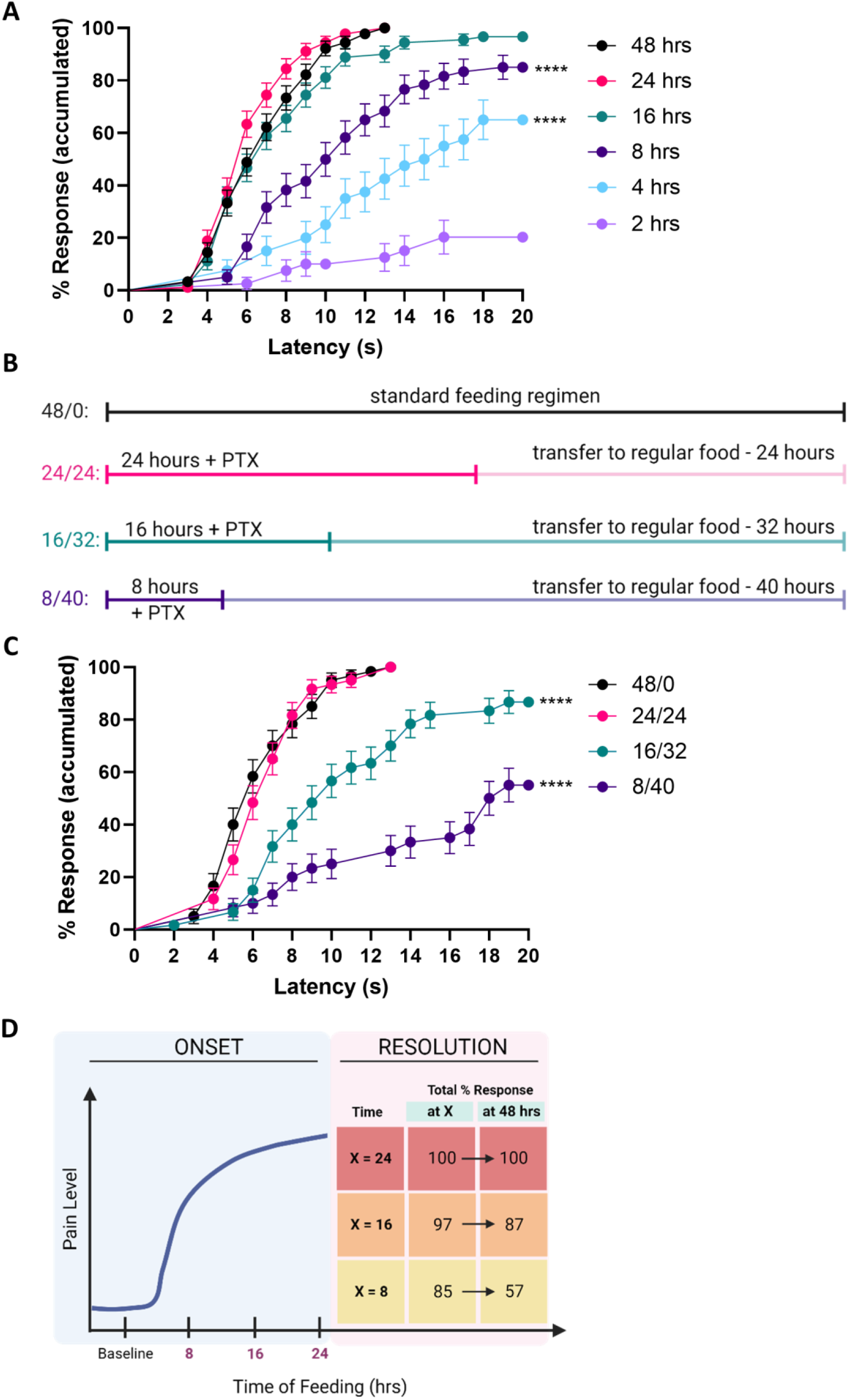
PTX-induced hypersensitivity is rapid onset and does not resolve at the larval stage. A. Quantitation of thermal nociceptive hypersensitivity (38.5°C thermal probe) of control larvae fed 10 µM PTX for the indicated times and assayed immediately afterward. For statistical comparisons, all timepoints were compared to the 48 hrs positive control (n=60 larvae per timepoint, ****p<0.0001 (Log-Rank Mantel-Cox)). B. Schematic of feeding regimen used to study temporal resolution of PTX-induced hypersensitivity. Standard feeding regimen is 48 hours on PTX. To study resolution, this 48-hour interval was broken into shorter times of feeding followed by recovery on regular food (no PTX), as indicated. C. Quantitation of thermal nociceptive hypersensitivity (38.5°C thermal probe) of larvae fed 10 µM PTX for the indicated times followed by recovery period (as in Figure 2B) and assayed at the end of the 48-hour period. Statistically, all timepoints were compared to the 48/0 positive control. (n=60, ****p<0.0001 (Log-Rank Mantel-Cox)) D. Graphical summary of onset and resolution data of PTX-induced thermal hypersensitivity for larvae fed 10 µM PTX for all or a subset of a 48-hour feeding period.

To study the temporal resolution of hypersensitivity, we devised a modified feeding scheme (see schematic, Figure 2B) followed by testing thermal nociceptive behavior. This scheme decreased the initial amount of time larvae were fed on 10 µM PTX in order to allow for a recovery period on regular instant fly food within a total 48-hour feeding period. Larvae fed PTX for 24 hours and allowed to recover for 24 hours were not significantly different, behaviorally, from larvae fed PTX for 48 hours (Figure 2C). However, decreasing PTX feeding time and increasing the recovery period significantly reduced larval hypersensitivity (Figure 2C). As the initial feeding time decreased the percent of total responders did as well, with a concomitant increase in average response latency (Figure 2C). However, larvae fed on PTX-containing food for 8 hours (and allowed to recover on untreated food for 40 hours), were still not recovered to baseline (Figure 2C). Figure 2D schematizes the onset data and compares the total percent responders tested immediately after a given feeding period (as in Figure 2A) with that same feeding period followed by recovery (as in Figure 2C). Together this data shows that PTX-induced thermal hypersensitivity has a rapid onset, within 24 hours, and does not completely resolve within the larval period.

### PTX causes alterations in nociceptor dendritic morphology

UV-induced and diabetes-induced hypersensitivity are not accompanied by obvious morphological changes in nociceptive sensory neurons (Babcock *et al*., 2009; Im *et al*., 2018). However, neuronal damage to both central axonal projections and peripheral dendritic arbors (Brazill *et al*., 2018; Shin *et al*., 2021) caused by PTX has been reported in *Drosophila*. We found that PTX treatment of larvae for 24 hours at 1 and 5 µM leads to increased dendritic branching in class IV peripheral sensory neurons compared to controls (Figure 3, A-C). Close up views of this are shown in Figure 3, A′-C′. Sholl analysis showed that the number of intersections increased in a broad peak centered at roughly 200 microns from the soma compared to untreated controls (see quantitation in Figure 3, E-G). In larvae fed 10 µM PTX, neurons showed signs of degradation which is characterized by beading not localized to branch points (Figure 3D and quantitation in Figure 3H). Thus, PTX feeding results in two distinct morphological consequences-dendritic hypersprouting at low concentrations (1 and 5 µM) and apparent damage, indicated by beading, at 10 µM.

**Figure 3.**
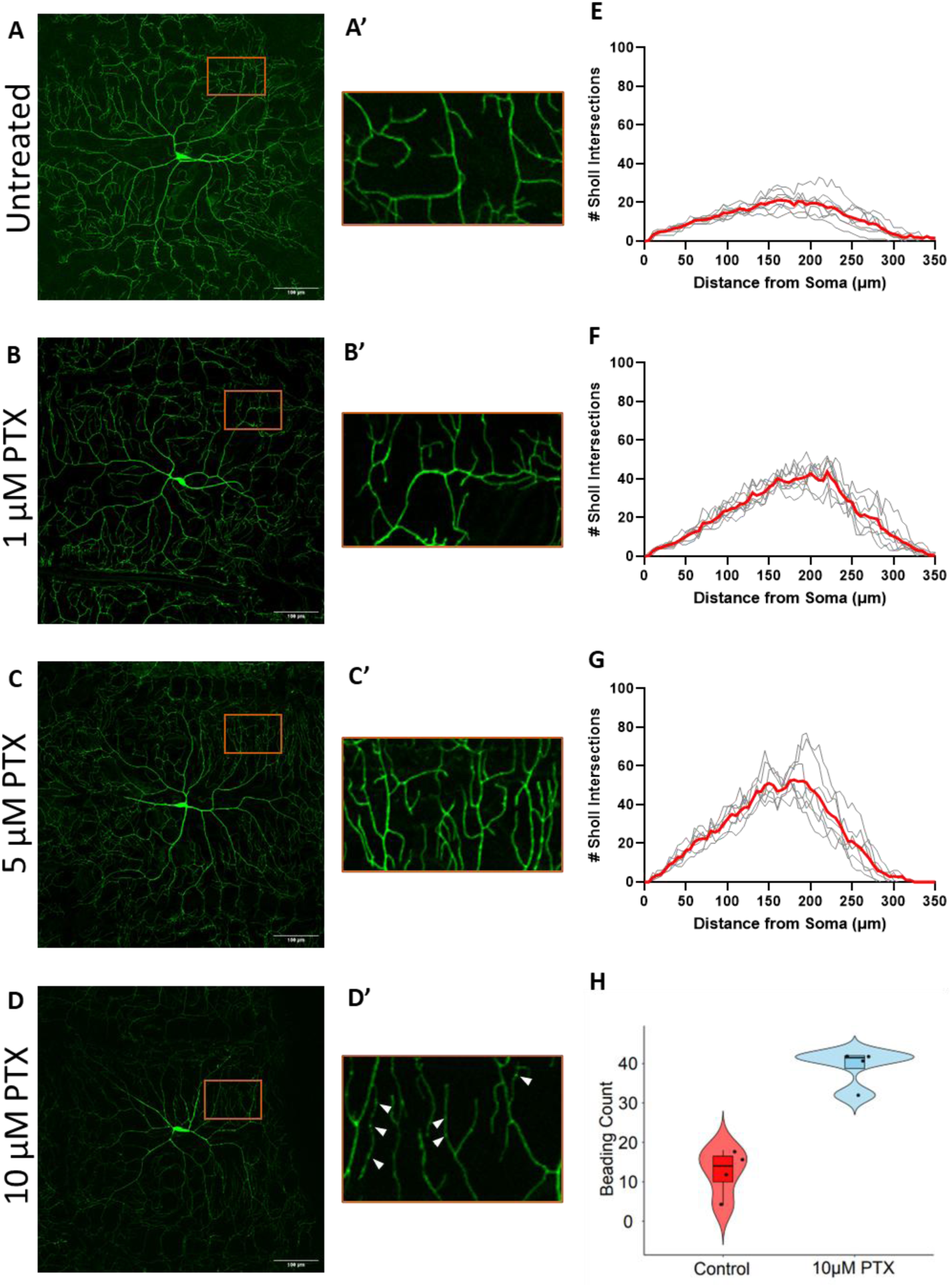
PTX causes alterations in thermal nociceptor dendritic morphology. A-D. Fluorescent confocal images of class IV peripheral sensory neurons in live larvae of the genotype *ppk1.9-gal4 > UAS-mCD8-GFP* larvae. Untreated (A) and fed 1 µM (B), 5 µM (C), and 10 µM (D) PTX. Images were taken at 20x magnification. A′-D′. Zoomed in images of dendrites from images 3A-D of the areas shown by the red boxes. Arrowheads in D′ indicate points of beading along the dendritic shafts. E-G. Quantitation of dendritic branching using Sholl analysis represented by number of Sholl intersections versus distance from the neuronal soma. Average of all measured neurons shown as red line. (n=7) H. Quantitation of dendritic degradation by counting points of beading (see methods for details) along dendritic shafts. (n=4 neurons per condition, ***p= 0.0005 (unpaired t-test)).

### Signaling pathways that regulate UV-induced thermal hypersensitivity are not required for PTX-induced hypersensitivity

To investigate the molecular mechanisms underlying PTX-induced hypersensitivity, we first tested signaling pathways known to mediate acute thermal hypersensitivity after injury. These include Tumor necrosis factor (TNF) signaling (Babcock *et al*., 2009), Tachykinin signaling (Im *et al*., 2015), and Hedgehog signaling (Babcock *et al*., 2009). As in these prior studies, we combined the nociceptive sensory neuron driver *ppk1.9-Gal4* with *UAS-RNAi* lines targeting essential components of each pathway to impair signaling in peripheral thermal nociceptors. Loss of function of these targeted genes: *Wengen* (a *Drosophila* TNF receptor) (Kanda *et al*., 2002), *Smoothened (Smo),* the signal transducer of the Hedgehog pathway) (van den Heuvel and Ingham, 1996), and the *Drosophila* Tachykinin receptor (*TkR99D*) (Monnier *et al*., 1992) did not alter thermal hypersensitivity compared to relevant Gal4- and UAS-alone controls when fed PTX (FigureS3). This suggests that these acute signaling pathways are not involved in PTX-induced hypersensitivity and that PTX-induced hypersensitivity is genetically distinct from UV-induced hypersensitivity.

### ILP4 and InR differentially regulate PTX-induced hypersensitivity

We next asked whether pathways that regulate persistence of injury-induced nociceptive sensitization also regulate PTX-induced hypersensitivity. Insulin-like signaling (ILS) in flies is required to attenuate acute nociceptive sensitization following injury and in diabetic-like states (Im *et al*., 2018). We first asked whether larvae mutant for ILPs had an effect on baseline nociception and found several that altered baseline thermal nociception. ILP5 and ILP3 mutants were slightly hypersensitive compared to controls while ILP4 and ILP6 exhibited decreased sensitivity (Table S1). Of these, ILP4 had the strongest phenotype and so we focused our analysis of PTX-induced hypersensitivity on this peptide and its putative receptor, InR. A full dose-response of temperature versus rolling behavior revealed that ILP4 mutant larvae exhibited a general shift towards less responsiveness across the full temperature range, with the exception of 38 °C-the low end of the curve (Figure S4; compare to curves in Figure S1).

To assess PTX-induced nociceptive changes, we tested ILP4 mutant larvae (Gronke *et al*., 2010; Colombani *et al*., 2012; Garelli *et al*., 2012) fed for 24 hours on 1, 5, and 10 µM PTX for thermal nociception and compared them to controls. ILP4 mutants were significantly less sensitive to a 38.5 °C thermal stimulus compared to controls when fed on 10 µM PTX (Figure 4A). Interestingly, they showed no difference from control larvae at lower PTX concentrations (1 and 5 µM) (Figure 4, B and C). We also investigated the involvement of InR, hypothesized to be the receptor for ILP4 (Brogiolo *et al*., 2001) in PTX-induced hypersensitivity. To test this, we expressed a *UAS-InR^RNAi^*line previously used to assess thermal nociception (Im *et al*., 2018) in class IV peripheral thermal nociceptors, using *ppk1.9-Gal4*. Compared to Gal4- and UAS-alone controls, expression of *UAS-InR^RNAi^* showed significantly increased sensitivity when fed on 10 µM or 5 µM PTX (Figure 4, D-F). These results on acute PTX-induced thermal hypersensitivity suggest that ILP4 (less hypersensitive at high PTX) and InR (more hypersensitive across several PTX concentrations), have different roles in regulating this process. This is surprising given that InR does seem to be a bona fide biochemical receptor for ILP4. Expressing InR in HEK-293 cells conferred the ability of these cells to bind to fluorescently labeled ILP4 and ILP5 (a known positive control ligand (Sajid *et al*., 2011) versus empty-vector transfected control cells (Figure S5).

**Figure 4.**
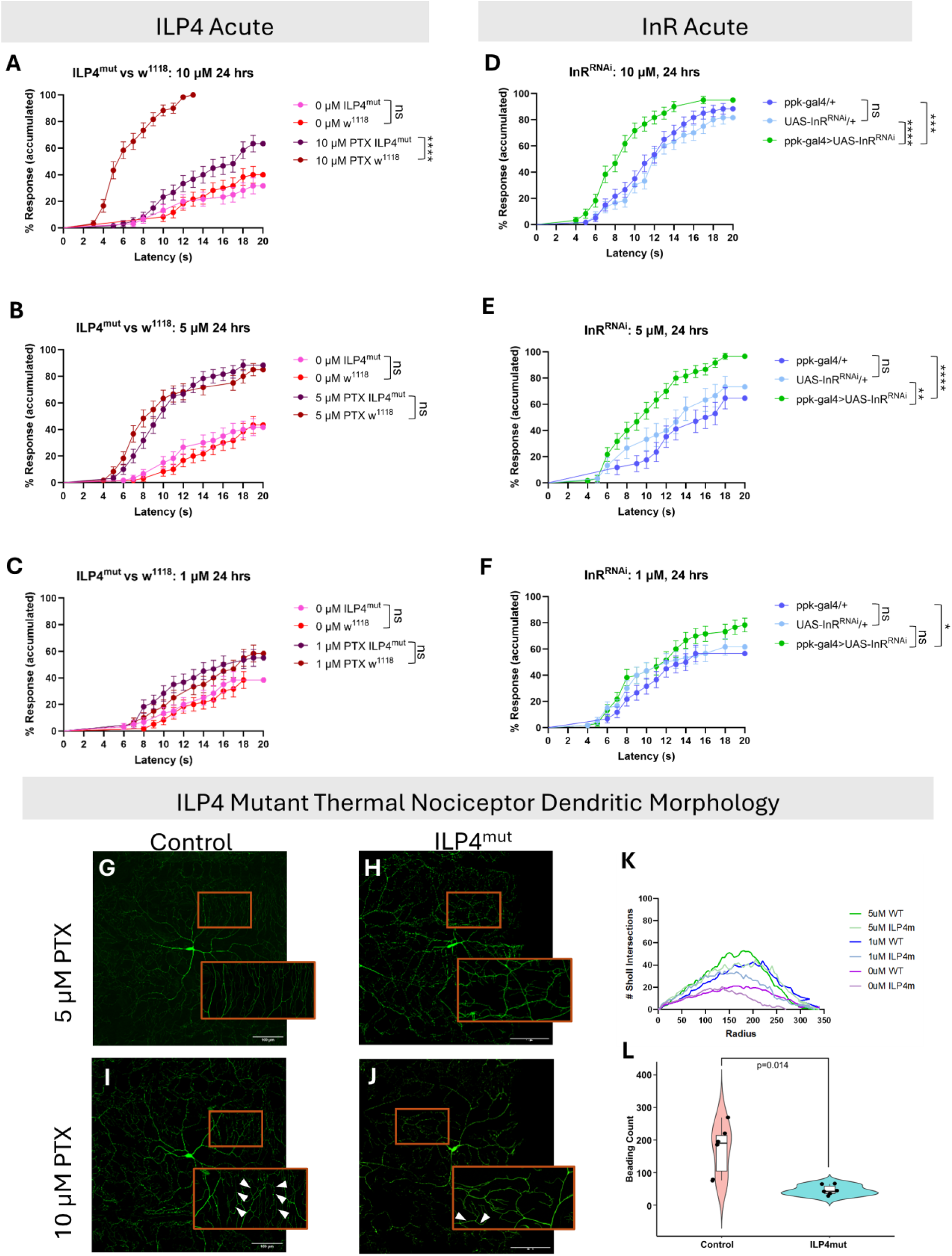
ILP4 and InR differentially regulate PTX-induced thermal hypersensitivity. A-C. Quantitation of thermal nociceptive hypersensitivity (38.5°C thermal probe) of *ILP4*-mutant larvae (dark purple) and w^1118^ control larvae (maroon) fed 10 µM (A), 5 µM (B) and 1 µM (C) PTX for 24 hours. Controls lacking PTX are shown in pink (*ILP4*) and red (*w^1118^*) respectively. Statistics, as indicated, compare untreated controls and treated groups to each other (Log-Rank Mantel-Cox). (n=60 per genotype, ns = not significant, ****p<0.0001). D-F. InR signaling-loss of function: Quantitation of thermal nociception (Heat probe = 38.5°C) for larvae expressing a *UAS-InR^RNAi^*transgene targeting the Insulin Receptor in class IV nociceptive sensory neurons (*ppk1.9-gal4*) and relevant Gal4-alone and UAS-alone controls fed 10 µM (D), 5 µM (E) and 1 µM (F) PTX for 24 hours (n=60 larvae per genotype). Stats: Log-rank Mantel-Cox, all comparisons versus the Gal4/UAS genotype (*p= 0.0193, **p= 0.0010, ***p= 0.0007, ****p<0001, ns = not significant). G-J Fluorescent livemount images of third instar larvae of the indicated genotypes and PTX concentrations. (G) *w^1118^* control, 5 µM PTX. (H) *ILP4^Mutant^*, 5 µM PTX. (I) *w^1118^* control, 10 µM PTX. (J) *ILP4^Mutant^*, 10 µM PTX. K-L Quantification of hypersprouting through Sholl analysis of control and *ILP4^mutant^* larvae (K) (n=6, each line is average for stated condition + genotype) and quantification of beading in control and *ILP4^mutant^* larvae (L). (n=6 neurons per condition, **p= 0.014 (unpaired t-test)).

Previous work showed that InR is required for the persistence of thermal hypersensitivity after injury and in diabetic-like states (Im *et al*., 2018). Thus, we also tested the persistence of PTX-induced hypersensitivity by feeding larvae of the relevant genotypes on PTX-containing food for 8 hours followed by recovery on normal food for 40 hours (as in Figure 2B). At all of the PTX concentrations tested (10-1 µM), *ppk1.9-Gal4 > UAS-InR^RNAi^* larvae were hypersensitive compared to Gal4-alone and UAS-alone controls. (Figure S5Z-A) indicating that, similar to diabetes-induced hypersensitivity (Im *et al*., 2018), InR regulates the resolution of PTX-induced hypersensitivity.

Finally, because ILP-4 has a specific requirement for PTX-induced hypersensitivity at 10 µM (but not 5 or 1 µM) and because these concentrations of drug induce different morphological effects in control larvae (see Figure 3), we tested whether PTX-induced morphological changes in ILP4 mutant larvae. Sholl analysis revealed that ILP4 mutants have slightly less complex class IV dendritic arbors than control larvae (Fig. 4 bottom panels) in the absence of PTX, a fact that might account for their slight decrease in baseline thermal nociceptive hypersensitivity (Figure S4). At the lower concentrations of PTX (5 and 1 µM) hypersprouting was observed in ILP4 mutants, similar to that seen in control larvae (Figure 4K). Interestingly, ILP4 mutants were protected from PTX-induced beading/damage at 10 µM PTX (Figure 4L), which might explain their reduced hypersensitivity at this particular concentration of drug.

### ILP4 expression and tissue-specific function

In order to determine which tissue(s) might express ILP4 peptide, we examined ILP4 expression using an *ILP4-Gal4* reporter where Gal4 expression is controlled by ILP4 upstream regulatory sequences (Min *et al*., 2016). Crossing *ILP4-Gal4* to flies bearing a *UAS-DsRed2Nuc* (Lesch *et al*., 2010) transgene revealed tissues expressing this peptide (Fig 5A-D). These included the dorsal/terminal chemosensory organ, the larval pharynx, the larval salivary gland, and tracheal cells. Notably, no expression in either the central nervous system (Figure 5E) or the peripheral nervous system was observed.

We next used *UAS-RNAi* lines (Dietzl *et al*., 2007; Ni *et al*., 2011) targeting ILP4 to determine in which tissue(s) functional ILP4 might be expressed. Four publicly available *UAS-ILP4^RNAi^* transgenes were crossed to the ubiquitously expressed *daughterless-Gal4 (Da-Gal4)* driver to determine whether any of the transgenes could phenocopy ILP4 mutants in dampening PTX-induced thermal nociceptive hypersensitivity (Fig. S7A). Several of these lines phenocopied ILP4 mutants, providing independent genetic evidence for the role of ILP4. In addition to our usual quantitation of the thermal nociceptive behavior plotting cumulative percent response versus latency to rolling (see Figure 1D for example) we also performed an analysis (see materials and methods) to quantify the average size of the “latency gap” between larvae of the same genotype +/- PTX (Figure S7B). By either of these measures the most potent *UAS-ILP4^RNAi^* line was BL12890. Ubiquitous expression of this transgene using *Da-Gal4* resulted in a profound on-target reduction of ILP4 message levels versus Gal4 alone and *UAS-ILP4^RNAi^*alone controls, close to the low level of message observed in ILP4 mutants (Figure S7C).

Finally, we crossed tissue-specific Gal4 drivers that drive strong expression in each of the ILP4-expressing tissues with the most potent *UAS-ILP4^RNAi^* transgene to determine the functional tissue source of ILP4. When *UAS-ILP4^RNAi^*was driven in tracheae using *Btl-Gal4* (Shiga *et al*., 1996)no amelioration of PTX-induced hypersensitivity was observed (Fig. 5F). Similarly, only a minor reduction of PTX-induced hypersensitivity was observed when the dorsal organ/terminal organ (DO/TO) driver *Gr66a-Gal4* (Dunipace *et al*., 2001) was used (Figure 5G). To examine function in the salivary gland we first compared the larval expression of two drivers, *Sage-Gal4* (Fox *et al*., 2013) and *Fkh-Gal4* (Henderson and Andrew, 2000) (see figure S8). Both expressed strongly in salivary glands, but *Sage-Gal4* was used here as it had the least expression in other tissues. Expression of *UAS-ILP4^RNAi^*using *Sage-Gal4* strongly curtailed PTX-induced thermal nociceptive hypersensitivity versus Gal4-alone and UAS-alone controls (Figure 5H) suggesting that the larval salivary gland is the primary tissue source of functional ILP4 peptide. To test this idea further we performed a rescue experiment, expressing *UAS-ILP4* in salivary gland via either *ILP4-Gal4* (all ILP4-expressing tissues) or *Sage-Gal4* (salivary gland) in a homozygous ILP4 mutant background. Restoring ILP4 expression to all ILP4-expressing tissues (Figure 5K) or only in the salivary gland (Figure 5L) also restored a normal hypersensitivity response to PTX, providing further genetic evidence that salivary gland-derived ILP4 is required for a normal hypersensitivity response to PTX.

**Figure 5.**
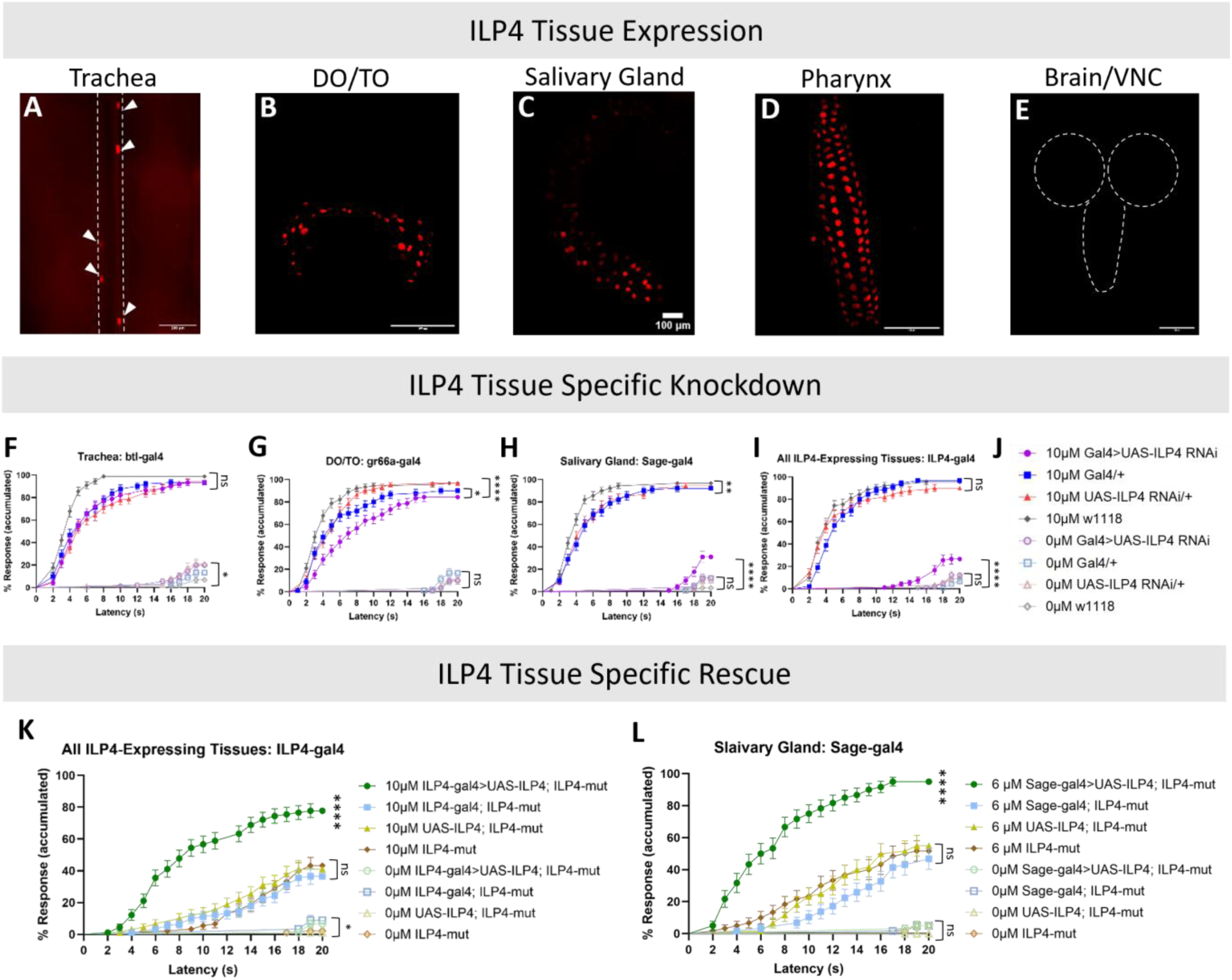
Expression and tissue source of functional ILP4. A-E Fluorescent confocal live wholemounts (A, B, D, and E) or dissected wholemount (C) images of tissues in larvae of the genotypes ILP4-Gal4>UAS-DsRed2Nuc. (A) Tracheal branches. (B) Dorsal/Terminal (DO/TO) organ. (C) Salivary gland. (D) Pharynx. (E) Brain/ventral nerve cord. F-J Quantification of thermal nociceptive behavior in larvae with the indicated tissue-specific Gal4 drivers driving *UAS-ILP4^RNAi^*. (F) Tracheae: *Btl-Gal4>UAS-ILP4^RNAi^*. (G) DO/TO: *Gr66a-Gal4> UAS-ILP4^RNAi^*. (H) Salivary gland: *Sage-Gal4>UAS-ILP4^RNAi^*. (I) All ILP4-expressing tissues: *ILP4-Gal4>UAS-ILP4^RNAi^*. (J) Legend for panels F-I. Stats: Log-Rank Mantel-Cox, n=90 per genotype, ns= not significant, *p=0.0216 (F) *p=0.0117 (G), **p=0. 0037, ****p=<0.0001. K-L Quantification of thermal nociceptive behavior in larvae of the indicated rescue genotypes and relevant controls. (K) *ILP4-Gal4* driving *UAS-ILP4* in all ILP4-expressing tissues in an *ILP4^mutant^*background and relevant controls. This experiment was performed with our original batch of PTX that was potent at 10 µM without adversely affecting survival. (L) *Sage-Gal4* driving *UAS-ILP4* in salivary gland in an *ILP4^mutant^*background and relevant controls. This experiment was performed with a new batch of PTX that was potent at 6 µM-see methods-concentrations higher than this adversely affected survival. Stats: Log-Rank Mantel-Cox, n=90 per genotype (K) and n=60 per genotype (L), ns= not significant, *p=0.0445, ****p=<0.0001.

## Discussion

Prior studies of PTX-induced hypersensitivity in *Drosophila* larvae focused mostly on axonal and dendritic damage to nociceptive sensory neurons (Bhattacharya *et al*., 2012; Brazill *et al*., 2018; Shin *et al*., 2021) and neuronal hypersensitivity at relatively high concentrations of PTX (Brazill *et al*., 2018; Shin *et al*., 2021). Here, we further explored the effects of PTX at lower concentrations. First, we showed that PTX induces thermal hypersensitivity at high concentrations that induce neuronal damage, as previously shown (Brazill *et al*., 2018). Surprisingly, we also showed that PTX-induced hypersensitivity can occur at concentrations that are up to 40 times lower than previously examined, including concentrations that do not induce morphologically obvious dendritic beading (see Figure 6 for a model that incorporates our results with respect to survival, morphology, PTX-concentration tested, behavior, and genetics). At high concentrations, PTX-induced hypersensitivity arises quickly, within 4 hours of feeding. This is consistent with the clinical observation that PTX-induced pain can arise as quickly as the first dose (Loprinzi *et al*., 2011). Also consistent with clinical data is the observation (Tanabe *et al*., 2013; Klein and Lehmann, 2021) that PTX-induced hypersensitivity is much slower to resolve. The fact that resolution is incomplete at the larval stage provides a potential platform for future study of the transition from acute to chronic hypersensitivity either in long-lived larvae (Wang *et al*., 2021) or at the adult stage.

**Figure 6.**
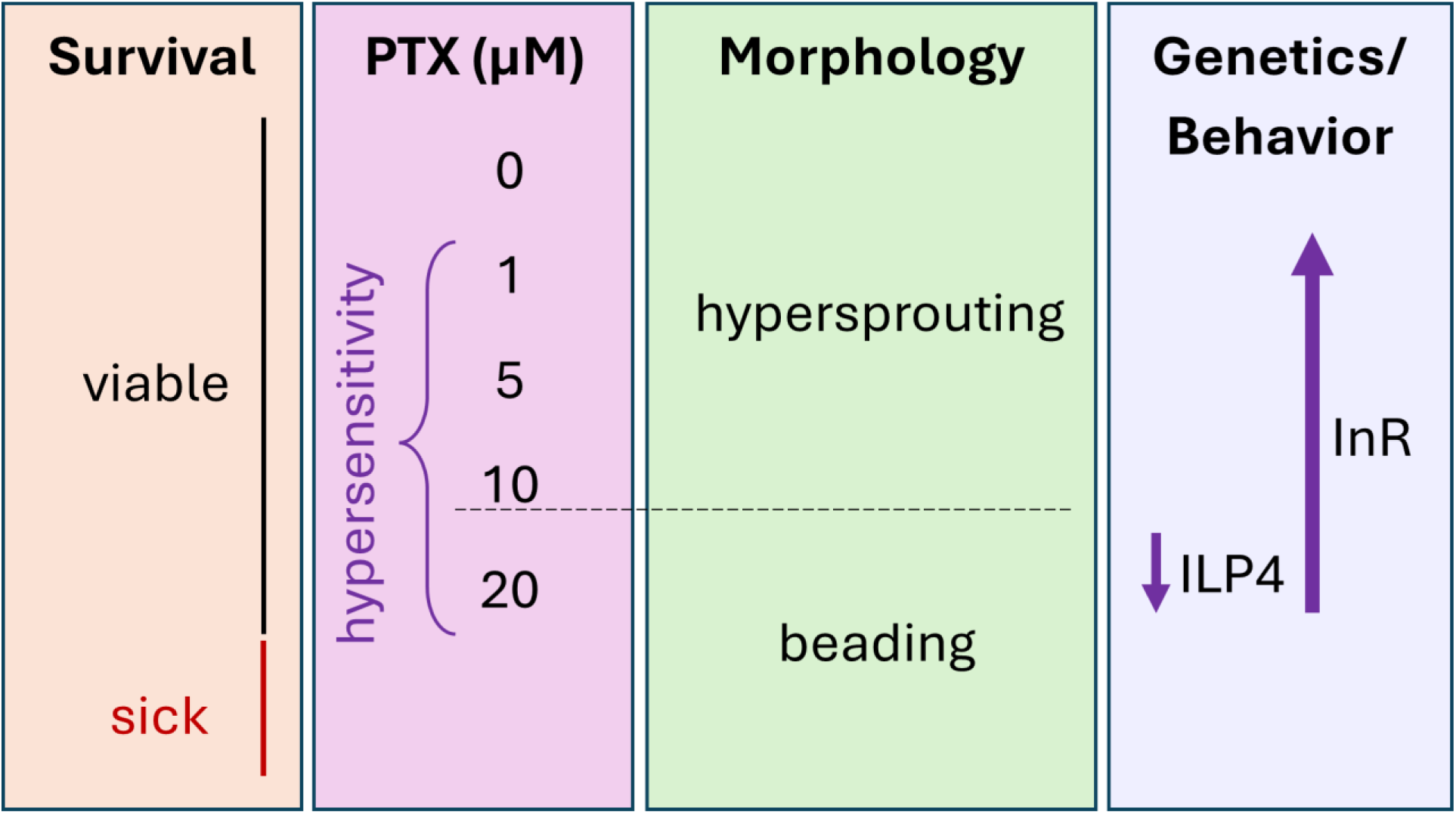
Model. Schematic summary of morphology, survival, behavior, PTX concentrations, and genetic results.

The relationship between PTX-induced hypersensitivity and morphological changes or damage to sensory neurons has long been debated. Sholl analysis of the morphology of class IV peripheral sensory neurons show shows two distinct PTX-induced morphologies not seen in controls. At lower PTX concentrations (1 and 5 µM), neurons exhibit increased dendritic branching, or “hypersprouting”. The branches affected are primarily the tertiary dendrites located over 150 microns from the neuronal soma. A previous study of mutants of the microtubule-severing protein Katanin (Stewart *et al*., 2012) showed that decreasing microtubule stability decreased both nociceptive responses and the stability of dendritic sprouts. Our results with PTX, a microtubule stabilizer, are exactly the opposite and suggest that the behavioral effects seen here are likely through direct or secondary effects on microtubules. At higher concentrations of 10 µM PTX and above (Bhattacharya *et al*., 2012; Brazill *et al*., 2018; Kim *et al*., 2020; Shin *et al*., 2021), we observed dendritic degradation characterized by beading on the dendritic shafts. Because thermal hypersensitivity is observed across the full range of non-toxic PTX concentrations, these two distinct morphologies raise the possibility that two separate mechanisms may be involved in PTX-induced hypersensitivity-one for low concentrations that show hypersprouting and another for higher concentrations that show beading (see Figure 6). Indeed, our genetic data, discussed below, strongly suggests that low- and high-PTX hypersensitivity are genetically distinct.

Prior studies of PTX-induced hypersensitivity in flies have indicated a neuroprotective role for NMNAT (Brazill *et al*., 2018) and the mitochondrial regulator PINK1 upon PTX exposure. Another reasonable hypothesis for PTX-induced hypersensitivity is that PTX treatment of larvae causes tissue injury that releases signals known to cause nociceptive hypersensitivity. Previous studies have shown that the TNF (Babcock *et al*., 2009), Tachykinin (Im *et al*., 2015), and Hedgehog signaling (Babcock *et al*., 2011) pathways are necessary for acute hypersensitivity following UV-induced damage to the barrier epidermis. Our data here suggest that these injury-induced pathways are not required for PTX-induced hypersensitivity.

We also explored the possibility that PTX-induced hypersensitivity was similar to diabetes-induced hypersensitivity by investigating insulin signaling (ILS) as a possible shared regulator. Partly this was motivated by a prior study showing that ILS regulates the persistence of injury-induced nociceptive sensitization in *Drosophila* larvae and the clinical observation of a similar loss of nerve fiber density in comparative studies of diabetic patients and patients receiving Paclitaxel (Lauria *et al*., 2003). It was also motivated by a survey of ILP mutants where we identified several that have modest alterations in baseline thermal nociception. Here, we found that ILP4 is required for full PTX-induced hypersensitivity in the acute timeframe (24 hours) at 10 µM PTX but not at 1 and 5 µM. Interestingly, this behavioral difference in ILP4 mutant larvae (required for PTX-induced hypersensitivity at 10 µM where beading occurs but not at 5 and 1 µM where hypersprouting occurs) is exactly mirrored by morphological analysis of ILP4 mutant larvae. In these mutants, dendritic beading on 10 µM PTX is suppressed, while hypersprouting at lower concentrations is not. The suppression of dendritic beading in ILP4 mutant larvae suggests that PTX-induced damage is an active process that requires ILP4 function.

Curiously, loss of InR causes increased hypersensitivity at all concentrations of PTX tested, a phenotype that is essentially opposite to that of ILP4. Biochemical and genetic evidence for InR binding has been previously shown only for two ILPs, ILP2 and ILP5 (Sajid *et al*., 2011; Park *et al*., 2014). Here, we confirmed that ILP5 binds InR and showed that ILP4 can bind this receptor as well, despite the genetic evidence that ILP4 is not likely acting through InR in this context. What is clear at this point is that ILP4 mutant larvae are protected from the degradative effects of high concentration PTX (10 µM) but not the hypersprouting effects of lower concentration PTX. A study in *C. elegans* found that a form of genetically-induced axonal degradation can be suppressed by either overexpression of NMNAT or by downregulation of ILS, suggesting that there may be a mechanistic connection between these pathways in neuronal protection (Calixto *et al*., 2012).

Analysis of an ILP4-Gal4 insertion where Gal4 is driven by ILP4 upstream regulatory sequences indicates that ILP4 is expressed in the dorsal/terminal chemosensory organ, the larval pharynx, and the salivary glands consistent with a potential role in feeding behavior (Wu *et al*., 2005). ILP4 is also expressed in tracheal cells but is curiously absent from neuronal tissues. The tissue that appears to produce functional ILP4 in the context of PTX-induced hypersensitivity is the salivary gland, since RNAi-mediated knockdown of ILP4 in this tissue phenocopies ILP4 mutants in protecting from PTX-mediated hypersensitivity. The suppression of PTX-induced hypersensitivity in ILP4 mutant larvae can be rescued by re-expressing ILP4 either from all ILP4-expressing tissues (using *ILP4-Gal4*) or only from salivary gland (using *Sage-Gal4*). Whether ILP4 expression is regulated by high concentrations of PTX is an interesting question for further analysis. Similarly, how salivary gland derived ILP4 acts to protect nociceptors from PTX-induced beading at high concentrations will be a topic for future investigation.

Presumably, ILP4 binds to a receptor other than InR to mediate protection from PTX-induced beading and hypersensitivity. At least one other ILP, ILP8, can bind a leucine-rich repeat-containing G protein-coupled receptor 3 (LGR3), a relaxin-like receptor (Vallejo *et al*., 2015). A study of avoidance of noxious light in *Drosophila* larvae, also a nociceptive response, implicated ILP7 in this escape behavior. Although InR was not analyzed in this study, an alternative receptor of the relaxin family, Lgr4, was. ILP7 was shown to bind Lgr4 and Lgr4 mutants had a similar defect in noxious light avoidance (Imambocus *et al*., 2022). These data suggest that LGR-family receptors (there are four in the fly genome) are attractive candidates for mediating ILP4’s regulation of PTX-induced hypersensitivity specifically at high acute concentrations (10 µM, 24 hours of feeding). A second interesting implication of our data is that there is likely an ILP (possibly ILP2 or ILP5) that acts directly through InR to mediate increased hypersensitivity in both the acute and persistent paradigms tested here. Further studies are needed to establish the binding and signaling activity of ILP4 and its role in PTX-induced hypersensitivity.

## Acknowledgements

We thank Bloomington, Korean, Kyoto, and VDRC stock centers for the indicated stocks. We thank Deborah Andrew and Stefan Luschnig for stocks. We thank Dr. Adriana Paulucci-Holthauzen at the Genetics Microscopy Core for providing technical support and assistance with imaging (NIH Instrumentation Grant S10 OD024976). This study was supported by NIGMS R35GM126929 (to MJG) and the American Legion Auxiliary Fellowship in Cancer Research from GSBS (to SS). Galko laboratory members read and commented on the manuscript.

## Abbreviations

(PTX): Paclitaxel
(ILP4): insulin-like peptide 4
(InR): insulin receptor
(ILS): insulin like signaling

