## Supplemental Data for "ILP4 and InR regulate Paclitaxel-induced hypersensitivity differently in *Drosophila* larvae"

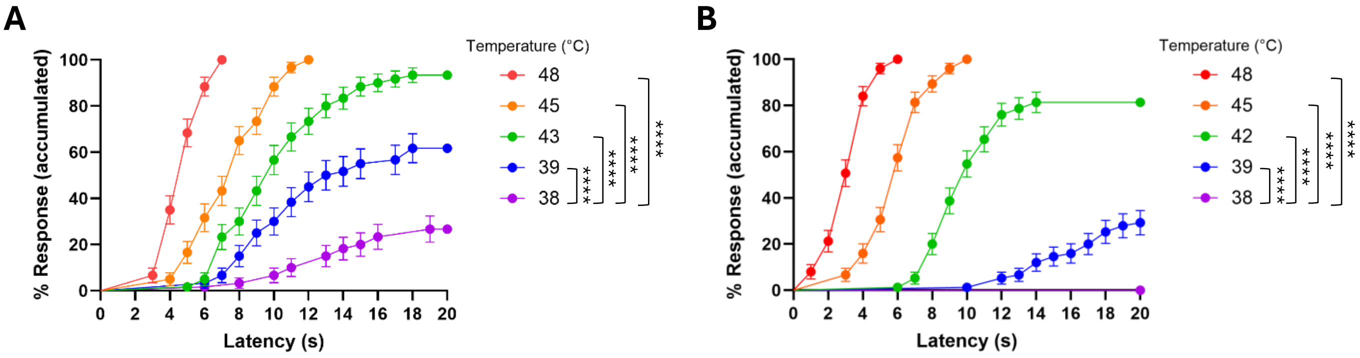


**Figure S1**. **Assay parameters for this study**.

Dose response quantitation of baseline thermal nociception with the particular heat probes used and users who performed behavioral experiments in this study. Each curve shows larval ($w^{1118}$) response at the indicated temperatures. Statistics (Log-Rank Mantel-Cox) compare each temperature to the next closest one.

A Curve from Sreepradha Sridharan (****p<0.0001, n=60).

B Curve from Yogesh Srivastava (****p<0.0001, n=30).

A

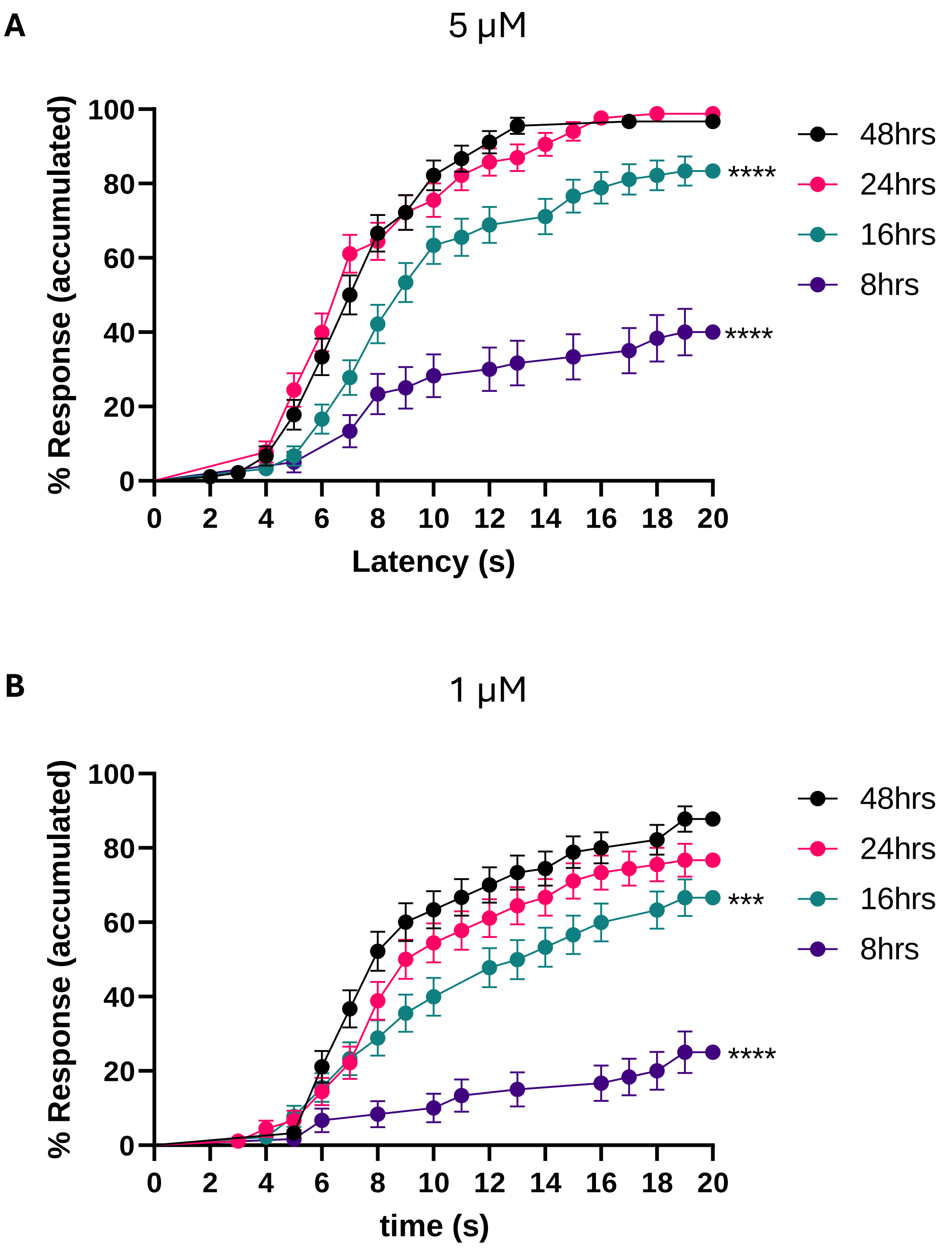


**Figure S2. Onset kinetics at lower concentrations of PTX.**

A. Quantitation of thermal nociceptive hypersensitivity (38.5°C thermal probe) of larvae fed 5 µM PTX for the indicated times and assayed immediately afterward. All timepoints were compared to 48 hrs positive control (n=60 larvae per timepoint, ****$p<0.0001$(Log-Rank Mantel-Cox)).

B. Quantitation of thermal nociceptive hypersensitivity (38.5°C thermal probe) of larvae fed 1 μM PTX for the indicated times and assayed immediately afterward. All timepoints were compared to 48 hrs positive control (n=60 larvae per timepoint, ****$p<0.0001$, ***$p= 0.0003$ (Log-Rank Mantel-Cox)).

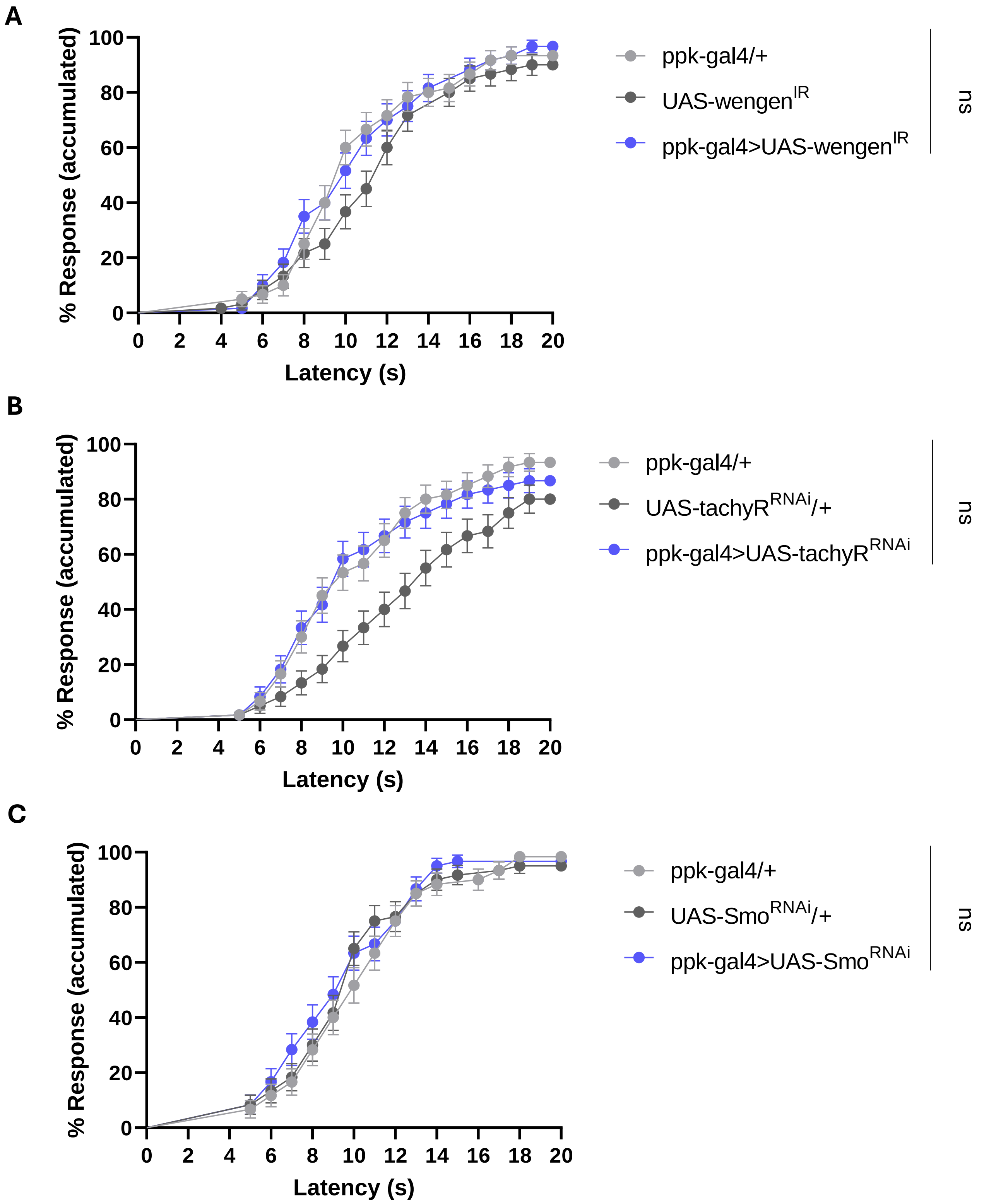
A
% Response (accumulated)
Latency (s)
ppk-gal4/+
UAS-wengen$^{IR}$
ppk-gal4>UAS-wengen$^{IR}$
ns
B
% Response (accumulated)
Latency (s)
ppk-gal4/+
UAS-tachyR$^{RNAi}$/+
ppk-gal4>UAS-tachyR$^{RNAi}$
ns
C
% Response (accumulated)
Latency (s)
ppk-gal4/+
UAS-Smo$^{RNAi}$/+
ppk-gal4>UAS-Smo$^{RNAi}$
ns

**Figure S3. Pathways known to regulate injury-induced thermal nociceptive hypersensitivity are not required for PTX-induced hypersensitivity.**

A. TNF signaling: Quantitation of thermal nociceptive hypersensitivity (Heat probe = 38.5°C) for larvae expressing a *UAS-RNAi* transgene targeting a TNFR (*UAS-TNFR$^{IR}$*) in class IV nociceptive sensory neurons (*ppk1.9-gal4*) and relevant controls (n=60 larvae per genotype). Stats: Log-rank Mantel-Cox, all comparisons versus the Gal4/UAS genotype not significant).

B. DTkR signaling: Quantitation of thermal nociceptive hypersensitivity (Heat probe = 38.5°C) for larvae expressing a *UAS-RNAi* transgene targeting DTkR (*UAS-TkR99D*) in class IV nociceptive sensory neurons (*ppk1.9-gal4*) and relevant controls (n=60 larvae per genotype). Stats: Log-rank Mantel-Cox, all comparisons versus the Gal4/UAS genotype not significant).

C. Smo signaling: Quantitation of thermal nociception (Heat probe = 38.5°C) for larvae expressing a *UAS-RNAi* transgene targeting the Hedgehog signal transducer Smoothened (*UAS-Smo$^{IR}$*) in class IV nociceptive sensory neurons (*ppk1.9-gal4*) and relevant controls (n=60 larvae per genotype). Stats: Log-rank Mantel-Cox, all comparisons versus the Gal4/UAS genotype not significant).

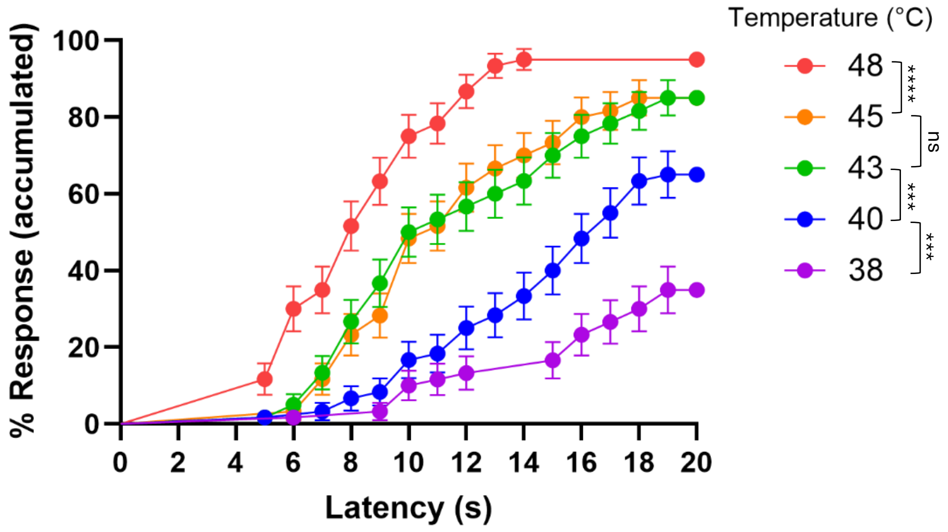


**Figure S4. Baseline thermal nociceptive dose-response of ILP4 mutant larvae**

Dose response quantitation of ILP4-mutant larvae. Each curve shows larval response at the indicated temperatures. Statistics (Log-Rank Mantel-Cox) compare each temperature to the next closest one. (****p<0.0001, 43°C vs 40°C ***p= 0.0004, 40°C vs 38°C ***p = 0.0007, n=60)

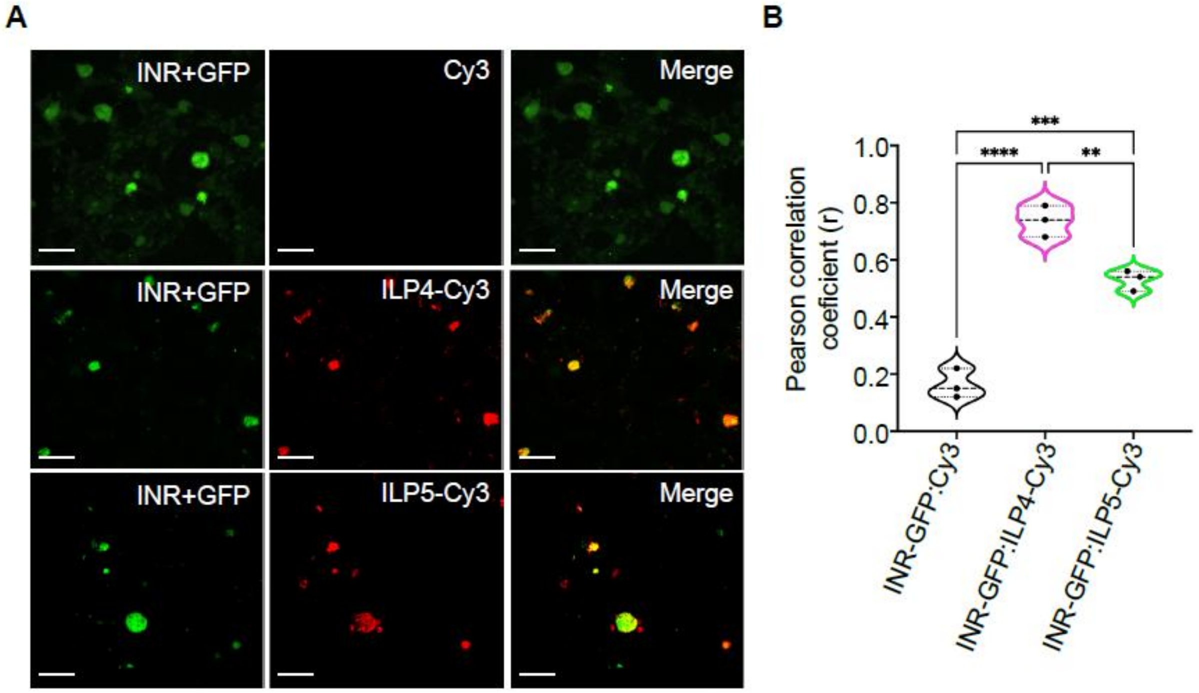


**Figure S5. Binding of ILP4 and ILP5 to InR.**

A. Fluorescent images of HEK-293T cells transfected with the indicated vectors (either pcNDA/FRT-GFP alone as control or this plasmid plus pcDNA-InR as experimental condition) and treated with the indicated fluorescent molecules to test binding (unconjugated Cy3; ILP4-Cy3; or ILP5-Cy3- see methods for details.

B. Quantitation of coincidence of green labeling (plasmid transfection) with red signal from Cy3 or labeled peptide binding.

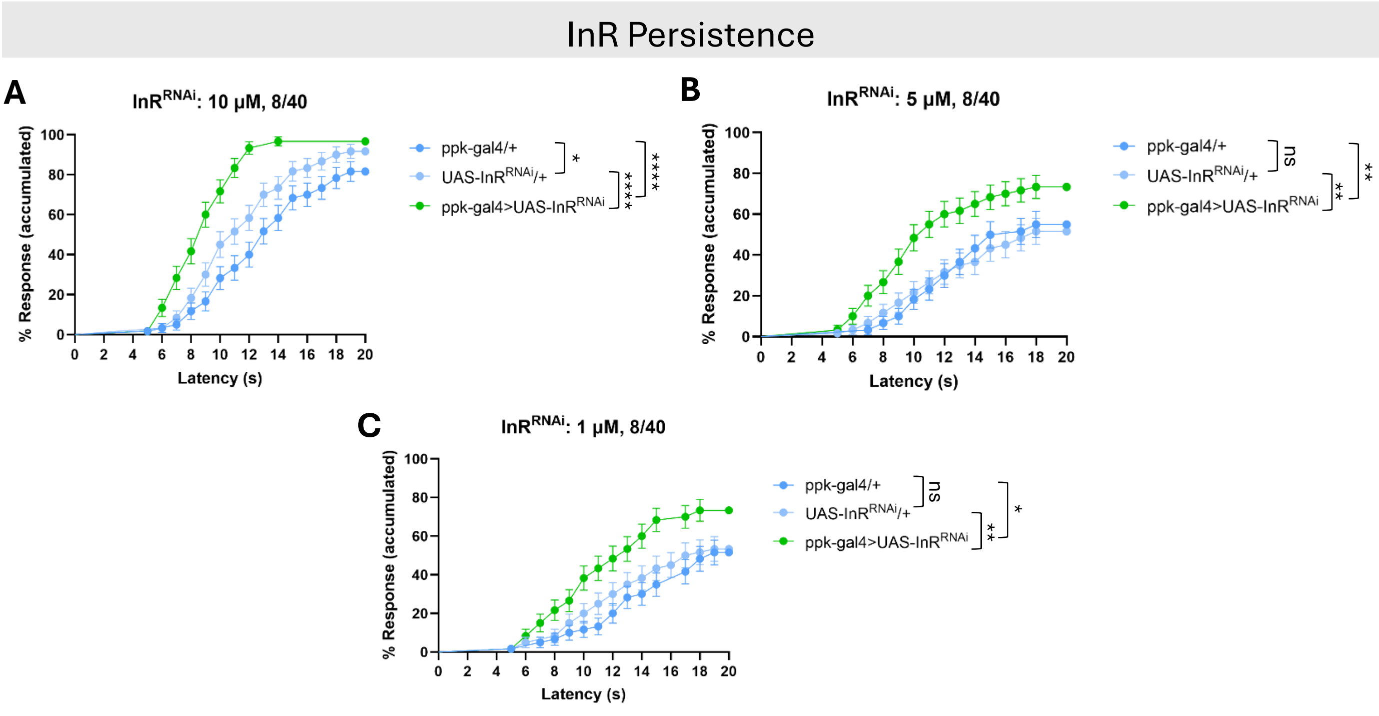


**Figure S6. Persistence of PTX-induced thermal nociceptive hypersensitivity.**

A-C. Quantitation of thermal nociceptive hypersensitivity (38.5°C thermal probe) of larvae fed 10 µM (A), 5 µM (B) and 1 µM (C) PTX for 8 hours followed by recovery period on normal food of 40 hours. Larvae expressing *UAS-InR*$^{RNAi}$ in class IV sensory neurons (*ppk1.9-gal4*) shown in green, and relevant Gal4-alone and UAS-alone controls (shades of blue) were assayed at the end of the 40-hour recovery period. (n = 60 larvae per genotype). Stats: Log-rank Mantel-Cox, all comparisons versus the Gal4/UAS genotype (A)*p= 0.0251, ****p<0.0001, (B) **p= 0.0034, **p= 0.0029, ns= not significant, (C) *p= 0.0107, **p= 0.0014, ns= not significant.

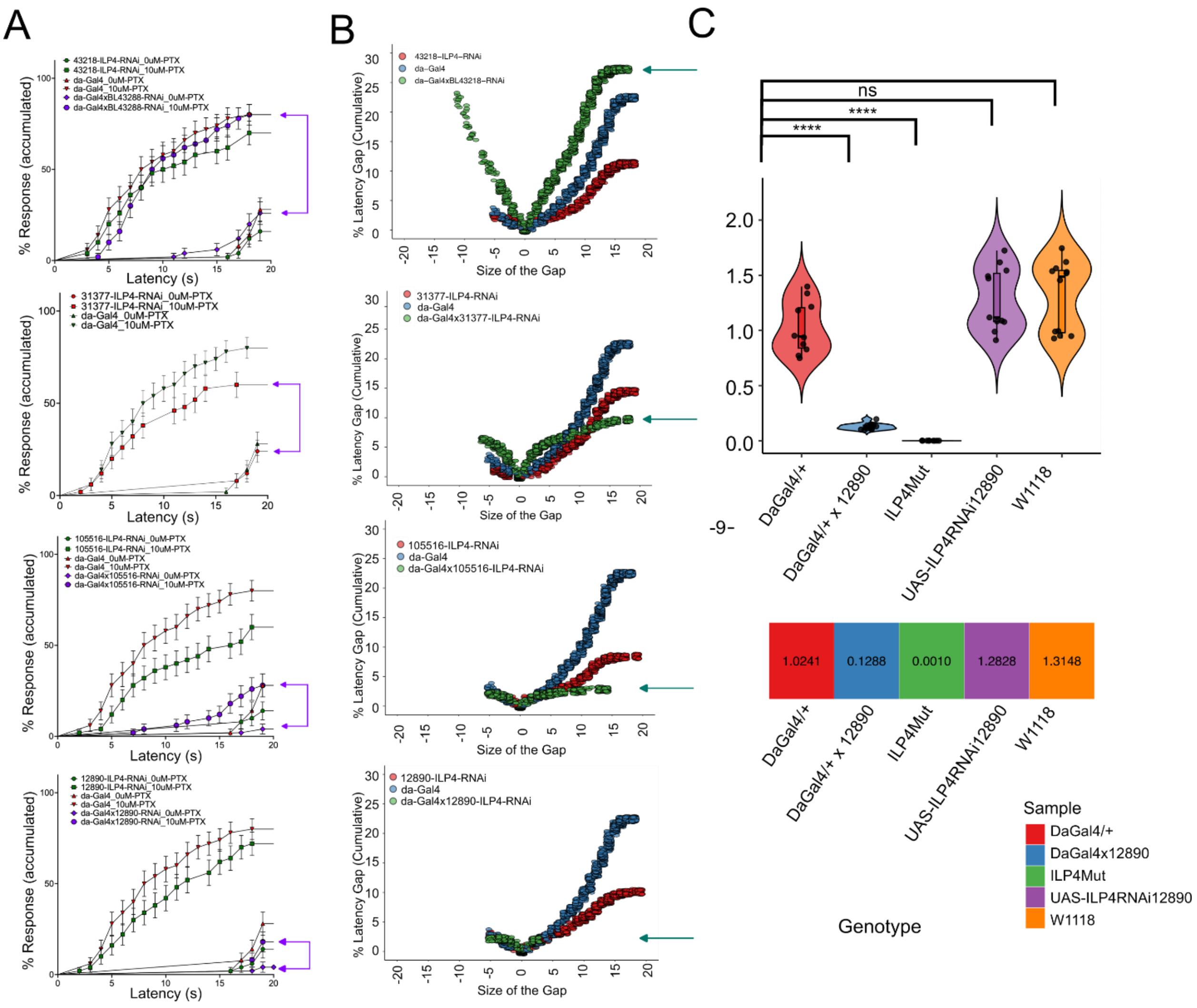




**Figure S7. Efficacy of *UAS-ILP4$^{RNAi}$* lines.**

A. Map of the four lines versus the ILP4 locus. Crib this from Flybase.

B. Quantification of PTX-induced thermal nociceptive hypersensitivity for each *UAS-ILP4$^{RNAi}$* line tested. Each graph contains data for the indicated RNAi line plus associated controls (Gal4 alone, UAS alone) performed in parallel +/- 10 μM PTX.

C. Quantitation of latency gaps (+/- PTX) for each *UAS-ILP4$^{RNAi}$* line and associated controls. See methods for details of the quantification.

D. qRT-PCR of the most efficacious RNAi line (BL12890) versus *ILP4*$^{mutant}$ and relevant genetic controls. Heat map indicates relative expression.

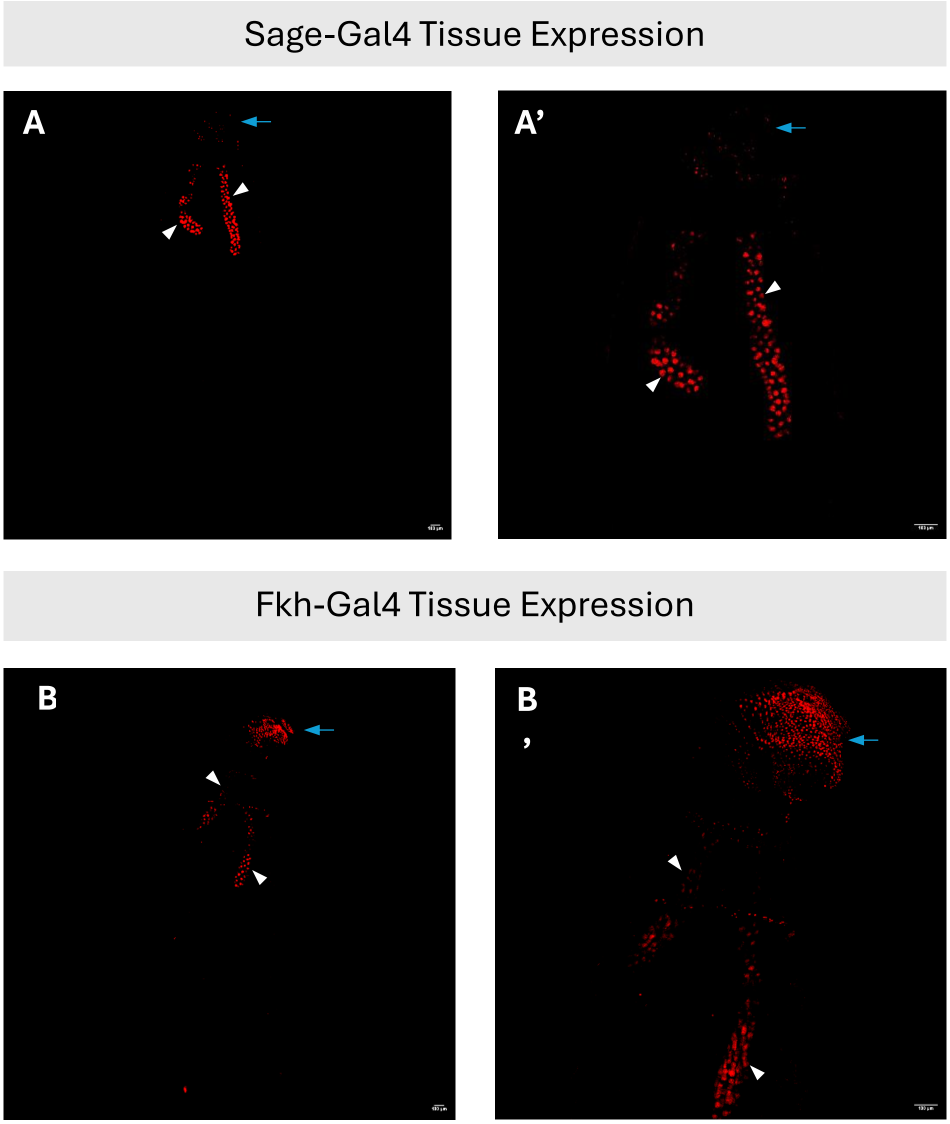


**Figure S8. Salivary gland Gal4 line comparison.**

A, A’. Fluorescent confocal wholemount images of tissues in larvae of the genotype Sage-Gal4>UAS- DsRed2Nuc. Whole body view (A) and close-up on salivary glands (A’) shown. White arrowheads denote salivary glands. Yellow arrow denotes DO/TO.

B, B’. Fluorescent confocal wholemount images of tissues in larvae of the genotype Fkh-Gal4>UAS- DsRed2Nuc. Whole body view (B) and close-up on salivary glands (B’) shown. White arrowheads denote salivary glands. Blue arrow denotes DO/TO.

| | **Thermal (48 °C)** |
|---|---|
| *ILP2* | NS |
| *ILP3* | ** |
| *ILP4* | **** |
| *ILP5* | *** |
| *ILP6* | * |
| *ILP7* | NS |
| *ILP8* | NS |

**Table S1. Table of ILP baseline thermal nociception phenotypes.**

The level of statistical difference of each ILP mutant versus $w^{1118}$ for thermal nociceptive hypersensitivity (48°C thermal probe) is shown. ILPs that showed increased sensitivity are shown in green, ILPs with decreased sensitivity are shown in blue (Log-rank Mantel-Cox test: ****p<0.0001, ***p= 0.0001, **p= 0.0043, *p= 0.0285, ns = not significant).

**Table S2. Genotypes of larvae used in each figure in this study.**

| Stock Genotype | Figure(s) Represented | Cat # | Acquired From |
|---|---|---|---|
| $w^{1118}$ | Figure 1, Figure 2, Figure 4 (A-C), Figure 5 (F-J), Figure S1, Figure S2 | - | Maintained in Galko Lab |
| ILP2-mutant | Table S1 | 30881 | BDSC |
| ILP3-mutant | Table S1 | 30882 | BDSC |
| ILP4-mutant | Figure 4 (A-C), Figure 5 (K-L), Figure S4, Table S1 | 30883 | BDSC |
| ILP5-mutant | Table S1 | 30884 | BDSC |
| ILP6-mutant | Table S1 | 30885 | BDSC |
| ILP7-mutant | Table S1 | 30887 | BDSC |
| ILP8-mutant | Table S1 | 33079 | BDSC |
| ILP4-gal4 | Figure 4(A-E, I-J) | 10011 | KDRC |
| UAS-ILP4 RNAi | Figure 4 (F-J), Figure S7 | 12890 | VDRC |
| UAS-ILP4 RNAi | Figure S7 | 105516 | VDRC |
| UAS-ILP4 RNAi | Figure S7 | 31377 | BDSC |
| UAS-ILP4 RNAi | Figure S7 | 43288 | BDSC |
| UAS-InR-ACT | Figure 4 (D-F) | 8263 | BDRC |
| UAS-tachykinin-RNAi | Figure S3 (B) | 103662 | VDRC |
| UAS-wengen-IR | Figure S3 (A) | 58994 | DGRC |
| UAS-Smo-RNAi | Figure S3 (C) | 9542 | VDRC |
| ppk1.9-gal4 (III) | Figure 3 | 32079 | BDRC |
| ppk-gal4 (II) | Figure 4 (D-F), Figure S3 (A-C) | 32078 | BDRC |
| da-gal4 | Figure S7 | - | BDRC |
| UAS-mCD8-GFP/CyO (II) | Figure 3 | 5137 | BDRC |
| ppk-gal4, UAS-mCD8-GFP/CyO ; ILP4-mutant/TM6B | Figure 4 (G-L) | - | Built in Galko Lab |
| UAS-dsNucRed11 (III) | Figure 5 (A-E), Figure S8 | - | Maintained in Galko Lab |
| Gr66a-gal4 (II) | Figure 5 (G) | 28801 | BDRC |
| btl-gal4 (II) | Figure 5 (F) | S273 | Dr. Stefan Lusching |
| Sage-gal4/CyO (II) | Figure 5 (H), Figure S8 | - | Deborah Andrew |
| Fkh-gal4 (III) | Figure S8 | - | Deborah Andrew |
| UAS-ILP4/CyO Act::GFP, ILP4-mutant/TM6B | Figure 5 (K-L) | - | Built in Galko Lab |
| ILP4-gal4/CyO Act::GFP, ILP4-mutant/TM6B | Figure 5 (K-L) | - | Built in Galko Lab |
| Sage-gal4/CyO Act::GFP, ILP4-mutant/TM6B | Figure 5 (K-L) | - | Built in Galko Lab |
